# CRISPRi perturbation screens and eQTLs provide complementary and distinct insights into GWAS target genes

**DOI:** 10.1101/2025.05.05.651929

**Authors:** Samuel Ghatan, Jasper Panten, Winona Oliveros, Neville E Sanjana, John A Morris, Tuuli Lappalainen

## Abstract

Most genetic variants associated with human traits and diseases lie in noncoding regions of the genome ^1^, and a key challenge is determining which genes they affect ^2,3^. A common approach has been to leverage associations between natural genetic variation and gene expression to identify expression quantitative trait loci (eQTLs) in the population ^4,5^. At the same time, a novel approach uses pooled CRISPR interference (CRISPRi) perturbations of noncoding loci with single-cell transcriptome sequencing ^6,7^. Here, we systematically harmonized and compared the results from these approaches across hundreds of genomic regions associated with blood cell traits. We find that while the two approaches sometimes identify the same target genes, there are considerable differences that affect biological inferences made from the data. CRISPRi preferentially maps highly proximal, constraint-enriched genes, whereas eQTLs recover multiple, often distal targets. By benchmarking against 1,075 gold-standard CRE–gene pairs linked to blood traits, we show that the two approaches identify largely distinct targets; when combined, they achieve a balance between accuracy and completeness of gene discovery. Our results offer guidance for improved design of CRISPRi and eQTL studies and highlight their joint potential as a powerful toolkit for interpreting disease-associated loci.

## Introduction

Genome-wide association studies (GWAS) have discovered hundreds of thousands of loci that associate with complex traits and diseases, but elucidating the functional mechanism of these variants remains challenging ^2,3^. Notably, ∼90% of GWAS-implicated variants lie in noncoding regions ^1^, making identifying causal genes difficult. Expression quantitative trait loci (eQTL) studies were the first genome-wide approach for associating variants with gene regulation. Nevertheless, despite their popularity, an emerging body of work highlights the limitations of relying exclusively on eQTL data to link GWAS variants to their putative target genes ^8,9^. Firstly, we still lack eQTL associations for many GWAS variants ^4,5^. When an eQTL does exist for a GWAS variant, it often finds genes unlikely to be causal to the trait ^4,8^, and its reliability in identifying the true causal gene is relatively poor compared to simple metrics such as assigning the closest gene as the causal gene ^10,11^. Recently, single-cell pooled CRISPR interference (CRISPRi) screens have emerged as a potentially powerful strategy for pinpointing causal genes at hundreds of GWAS loci ^6,7^. However, CRISPRi inhibits entire enhancers ^12^, producing effects that may not be analogous to single-nucleotide variation, and the studies are typically conducted in non-primary human cell lines, raising questions about the transferability of biological effects into primary cells and disease associations. The sensitivity or specificity of the CRISPRi approach for identifying likely causal GWAS genes has not been systematically evaluated, and the extent to which eQTL and CRISPRi approaches converge or diverge in identifying GWAS gene targets has been unknown.

### GWAS target gene discovery

We analyzed and fine-mapped GWAS loci from 44 blood trait studies from the UK Biobank and the Blood Cell Consortium (BCX) (data S1). To identify *cis*-eQTL target genes (eGenes) for GWAS variants, colocalization was performed with fine-mapped eQTL variants from 51 studies based on RNA-sequencing of whole blood, bulk RNA-seq of isolated blood cell types, and single cell RNA-sequencing (scRNA-seq) of peripheral blood mononuclear cells (PBMCs) (data S2) ^5,13–15^. To identify CRISPRi cis target genes (cGenes), we utilized and/or re-analyzed 12 CRISPRi studies targeting enhancers in K562 erythroleukemia cells (data S3) ^6,7,16^ and identified fine-mapped GWAS variants intersecting gRNA loci (Fig. 1a, mean PIP = 0.20). Overall, we identified 882 GWAS loci in cis-regulatory elements (GWAS CREs) that were targeted by a CRISPRi gRNA. Of these, 145 CREs (16%) had a cGene (FDR < 0.10, fig. S1a), and 578 (66%) had a colocalized eGene (PP.H4 > 0.5, fig. S1b) within 1 Mb (Supplementary Tables 4 & 5). In 104 CREs, both eGenes (n = 321) and cGenes (n = 130) were identified, with 81 overlapping CRE–gene target pairs shared between the two approaches (Fig. 1b).

**Figure 1.**
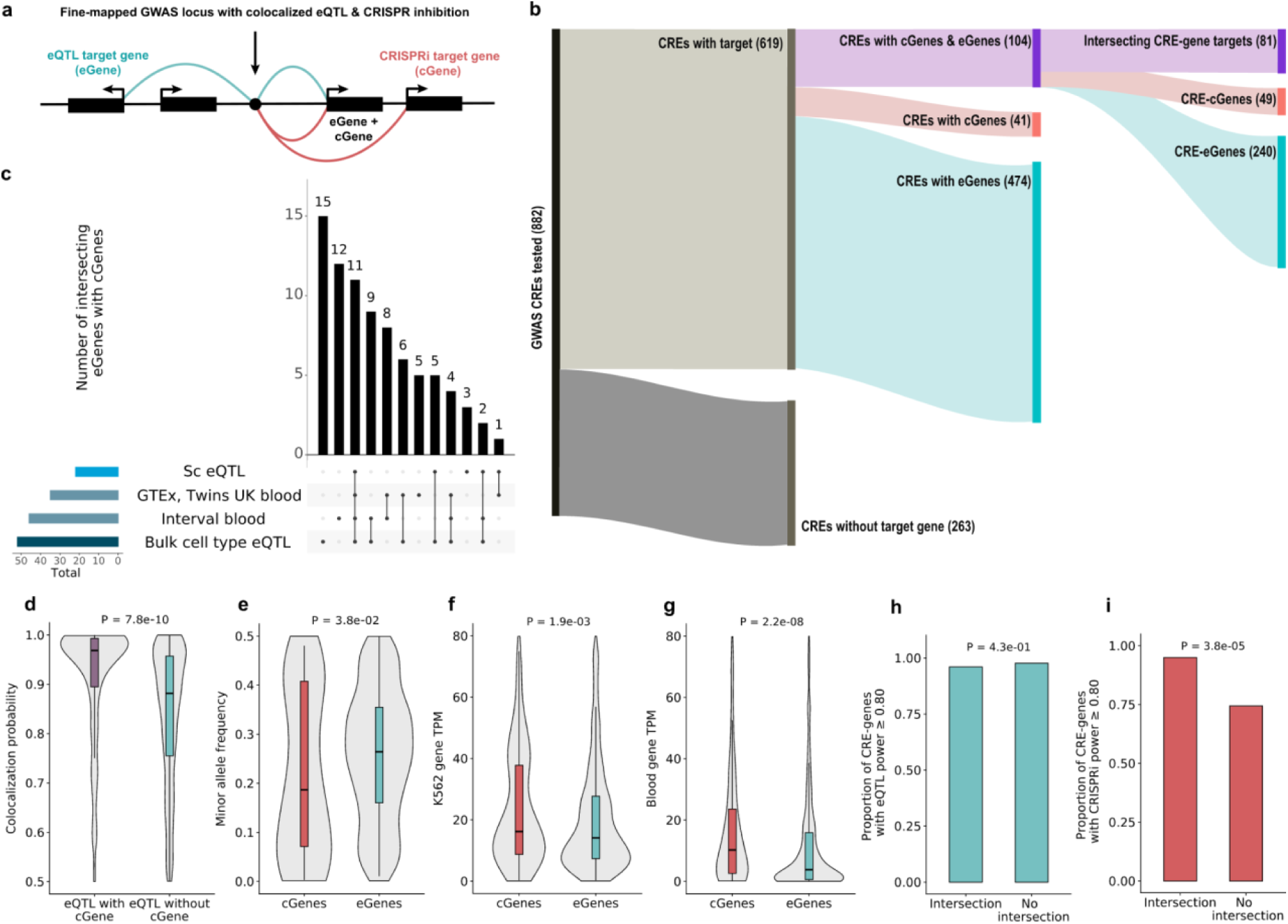
Discovery of eQTL and CRISPRi target genes for GWAS loci. **a**, Illustration of the comparison of gene targets for GWAS variants identified through eQTL and CRISPRi approaches. **b**, The number of CREs and genes discovered by eQTL and CRISPRi. **c**, The number of eGenes that intersect with cGenes across different eQTL study types. **d**, Comparison of GWAS–eQTL colocalization probabilities between eQTLs whose eGenes intersect a cGene and those that do not. **e**, Minor allele frequencies (MAFs) for GWAS variants uniquely mapped to eGenes or cGenes. **f-g**, Gene expression (transcripts per million; TPM) for cGenes and eGenes in K562 erythroleukemia cells **(f)** and in GTEx whole blood **(g)**. **h-i**, Simulated discovery power for eGene at aFC of 0.25 **(h)** and cGene at effect size 0.25 **(i)** between CRE-gene targets that intersect between CRISPRi and eQTL methods versus those that do not, at GWAS-associated CREs with both target types.

### CRISPRi and eQTL power shape the discovery of GWAS target genes

We first investigated how these discoveries were affected by various gene, CRE, and study features related to statistical power by analyzing patterns in the empirical data, complemented by simulations of both eQTL and CRISPRi discovery in each locus (Methods, Supplementary Tables 6 & 7, Extended Data Fig 2). Only 3 of the overlapping genes were uniquely identified in scRNA-seq eQTLs, pointing to limited power in current single-cell eQTL data in identifying new GWAS target genes, particularly for less abundant cell types (Fig. 1c, fig. S1c). Based on our simulations, the eQTL studies had a high mean power of 0.90 to detect associations with moderate effects of β = 0.25, but much larger samples would be needed to detect smaller effects (fig. S3c). While we observed additional GWAS colocalizations with eQTL sample size, this increase scaled sublinearly with sample size: INTERVAL is 8.3× larger than GTEx yet has only 2.7× more cis-eQTLs (56,959 vs 20,818) and 2.5× more colocalized loci (10,662 vs 4,292) (fig. S3d). Even though we analyzed only colocalized eQTLs (PP.H4 > 0.5), GWAS variants with eGenes that intersected cGenes had a higher colocalization probability (PP.H4) (*P* = 7.8 × 10^-10^, Fig. 1d), even after controlling for eQTL and GWAS P-values. These results suggest that when cell type isolation is technically feasible, bulk cell type eQTL studies hold advantages over sc-eQTL and bulk tissue studies in providing both good statistical power and cell type specificity, and that stricter criteria for eQTL colocalization may yield more reliable target inference.

Some of the differences between eGene and cGene discovery are simply due to cell type specific gene expression: in the 104 CREs with both cGenes and eGenes, 98% of cGenes were expressed in eQTL datasets, whereas only 76% of eGenes were expressed in K562 cells, underscoring the limitations of inferring causal genes from a single cell line. Correlation in rank concordance of trait associations was moderate (Spearman ρ = 0.58, P = 0.001; Fig. S4). Consistent with the erythroid lineage of K562, CRISPRi mapped genes for red-blood-cell GWAS loci more effectively (within-trait rank test, Δrank = rank_CRISPRi − rank_eQTL; median = −2.0; one-sided Wilcoxon P = 0.04), whereas eQTL studies tended to capture more genes for white-blood-cell traits (median = 5.0; P = 0.07). This highlights the importance of matching studied cell types to the studied GWAS traits.

cGenes were identified for GWAS variants at lower minor allele frequencies (MAFs) than eGenes (*P* = 0.04, Fig. 1e), a pattern recapitulated in simulations (fig. S3a). This is expected, as eQTL discovery power, similarly to other variant association studies, declines for low-frequency variants. In contrast, CRISPRi-based mapping is unaffected by allele frequency. Conversely, eGenes have significantly lower expression than cGenes in K562 cells (*P* = 1.9. × 10^−3^, Fig. 1f) and GTEx whole blood (*P* = 2.2 × 10^−8^, Fig. 1g), with simulations further indicating that CRISPRi discovery power is more affected by lower expression than eQTLs, likely deriving from sparsity of scRNA-seq data (fig. S3b). Altogether, eGene discovery power was similar for genes that intersect cGenes and for those that do not (*P* = 0.43, Fig. 1h), suggesting that the eQTL analysis is not lacking power to capture these target genes. The opposite was true for cGene discovery power, which was significantly higher for intersecting target genes compared to non-intersecting ones (*P* = 3.8 × 10^-5^, Fig. 1i), indicating that higher powered CRISPRi data would result in better alignment with eQTL results.

### Biological properties of eQTL and CRISPRi discoveries

Next, we examined the biological properties of the discovered eGenes and cGenes. While most cGenes (81%) and eGenes (68%) lie within 100 kb of the GWAS CRE (Fig. 2a), cGenes are more proximal: 41% are within 10 kb, compared to only 23% of eGenes. As a baseline, 21% of CREs without a cGene or eGene have the closest gene within 10 kb (Fig. 2b). The proximity of CRE–cGene pairs is notable given that CRISPRi screens specifically targeted regions more than 1 kb from transcription start sites, and repression by KRAB/MeCP2 is generally limited to local chromatin ^6,7,17^. Thus, we do not expect CRISPRi to cause gene repression by inhibition of gene promoters. eQTL mapping more often identifies genes beyond the 2nd closest gene (64% vs. 23% for cGene-linked CREs, *P* = 1.0 × 10⁻^10^, Fig. 2c). Additionally, CRISPRi identifies a smaller number of genes, with 82% of cGene-linked CREs and 33% of eGene-linked CREs being associated with a single gene (*P* = 2.0 × 10⁻^26^, Fig. 2d). This difference reflects not only the power of eQTL mapping but also the broader range of eQTL studies and cell types analyzed, with a median of two distinct eQTL studies identifying unique genes per CRE (fig. S5a). These results indicate that eQTL mapping better captures weaker regulatory effects that span larger genomic distances. This is important for identifying more distal target genes as more distal CRE-promoter interactions are, on average, weaker ^18^ (*P* = 7.5 × 10^-05^, fig. S5b) and thus harder to detect. Larger future CRISPRi studies with more diverse cell types might bring the two approaches closer to alignment.

**Figure 2.**
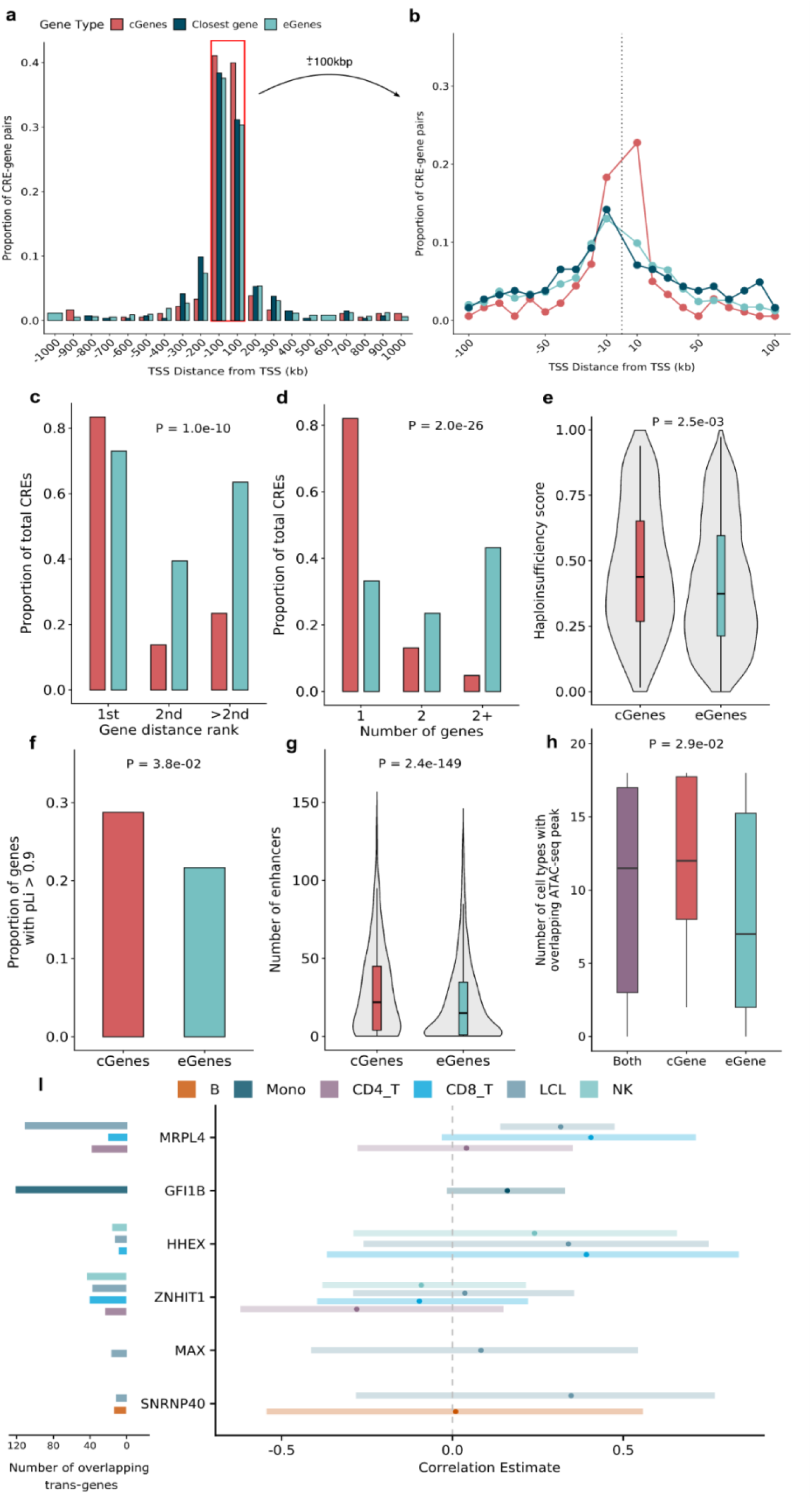
Biological properties of eQTL and CRISPRi discoveries. **a,** The distance between GWAS-associated CREs and target gene TSS, comparing cGenes, eGenes, and the closest gene to GWAS CREs where no significant association was identified, with **b** showing the region within 100 kb of TSS. **c,** The proportion of GWAS-associated CREs linked to their closest, second closest, or further away genes. CREs may be counted in multiple categories if they are associated with multiple genes. **d**, The proportion of GWAS-associated CREs linked to one, two, or more genes. **e**, Haploinsufficiency (predicted by Episcore) for cGenes and eGenes. **f**, The proportion of cGenes and eGenes with loss-of-function intolerance metric pLI > 0.9. **g**, The number of enhancers for cGenes and eGenes. **h**, Number of blood-cell types in which a CRE has open chromatin for CREs with both eGenes and cGenes, only cGenes, and only eGenes. **i**, Trans-gene analysis, with the y-axis indicating the cis-genes through which trans-effects occur. The left-hand side of the plot shows the number of overlapping trans eGenes and cGenes, and the correlation and its confidence interval of the effect sizes are shown on the right-hand side.

It has been proposed that, due to natural selection, eQTLs that rely on natural genetic variation are biased against selectively constrained effects ^19^. Indeed, we observed that cGenes were more haplosensitive, measured by Episcore predictions (*P* = 2.5 × 10^-3^, Fig. 2e), pHaplo scores (*P =* 0.01), and pLI scores ≥ 0.9 (P = 0.04, Fig. 2f). This suggests that CRISPRi is better at identifying genes that do not tolerate dosage variation. Furthermore, cGenes had a higher number of associated enhancers (*P* = 2.4 × 10^−149^, Fig. 2g) and super-enhancers (*P* = 3.4 × 10^−4^), suggesting a more complex regulatory architecture and tighter regulatory control than eGenes. These results highlight the potential of experimental perturbations such as CRISPRi to discover genes under selective and regulatory control, which are more likely to be relevant for diseases and traits ^20^. However, it remains unclear how often highly constrained genes undergo more buffering ^21^ to the extent that their expression changes are harder to detect even after CRISPR perturbations.

From ATAC-seq data spanning 18 blood cell types (Supplementary Methods), we observed that CREs with only eGenes had more cell-type specific open chromatin than those with also or exclusively cGenes (Fig. 2h; Kruskal-Wallis rank sum test, *P* = 0.03). This suggests that cell type differences between primary cell types in eQTL studies and the K562 line contribute to the sharing of detected regulatory effects, and that experiments in K562s tend to pick up regulation that is shared between many primary cell types. This is potentially due to K562 being a progenitor cell type, or because distal enhancers that are shared across cell types are also stronger on average, and thus we have more power to detect their associated genes. Notably, 12% of eGenes were located in regions of closed chromatin across all blood cell types (excluding K562), compared with only 5% of cGenes (P = 0.02), supporting the idea that CRISPRi-based mapping is especially sensitive to classical biochemical enhancer features ^6,7^. Hi-C data in K562 cells revealed that enhancer-gene targets from physical chromatin contacts were more likely to overlap with cGenes than eGenes (*P* = 1.8 × 10^-5^, fig. S5c), although 53% of the GWAS CREs tested had no annotated Hi-C contacts.

### Shared cis-gene anchors reveal variable *trans*-regulatory landscapes across datasets

We next examined how CRISPRi-identified *trans*-regulatory genes (*trans*-cGenes) compared with eQTL-based *trans* genes (*trans*-eGenes) for the same GWAS CREs acting through a shared *cis*-gene. We detected significant *trans*-cGene effects for 16 GWAS CREs (FDR < 0.10; 72–3,870 trans-cGenes per CRE; fig. S6a, Supplementary Tables 8 & 9). At two of these loci (*GFI1B* and *HHEX*), trans-cGenes were detected by both Morris et al. and Gasperini et al.^6,7^, with high correlation in effect sizes (*GFI1B*: r = 0.88, P < 2 × 10^-16^, n = 921; *HHEX*: r = 0.92, P = 9.6 × 10⁻⁸, n = 18, Extended Data Figs. 6b & c). To identify *trans*-eGenes for these loci, we analyzed *trans*-eQTLs (FDR < 0.05) from single-cell PBMC data from OneK1K ^15^ and a large meta-analysis of lymphoblastoid cell lines (LCLs) (n = 3,892) ^22^. We discovered *trans*-eGenes for 11 out of 16 CREs (data S10), of which 6 had overlapping *trans*-genes (fig. S6d). The correlation between trans-eGene and trans-cGene effects was modest for most loci and often variable between cell types (r = -0.28–0.41, Fig. 2i), and significant only for the CRE regulating *MRPL4* in LCLs (r = 0.32, FDR = 9.9 × 10^-3^, data S11). Additionally, the overlap of trans-cGenes and trans-eGenes was not significant for any CREs or cell types (fig. S6e, data S11). These data indicate the potential for robust detection of trans-regulatory networks with both approaches, but also the need for further work to fully understand the drivers of shared and distinct patterns across a larger number of loci.

### Detection of gold-standard GWAS genes by eQTL and CRISPRi

To evaluate the ability of each method to identify genes that are likely to be true causal genes for blood trait phenotypes, we constructed a “gold-standard” gene list of 421 protein-coding genes in the studied loci (data S12). This included genes identified through loss-of-function (LoF) burden tests for 29 blood cell traits in the UK Biobank ^23^ and a curated set of genes linked to Mendelian blood disorders (Methods). eQTL mapping identified 65 of these, and 23 were discovered as cGenes, corresponding to 11% (65/578) of all eGenes and 22% (23/107) of all cGenes in the loci containing gold-standard genes (Fig. 3a). Only 10 genes were detected by both approaches. When a cGene overlapped a gold-standard gene, it was the closest gene to the GWAS CRE in 74% of cases, compared to 54% for eGenes, indicating the further potential of eQTLs to identify more distal target genes that are more difficult to identify as putative targets of GWAS loci. Interestingly, gold-standard eGenes had a similar eQTL effect size distribution as other eGenes, indicating that even weak eQTL effects will pinpoint true target genes (Extended Data Fig 7) ^9,24^. However, the patterns of gene target discovery are highly variable between the loci (Fig. 3d, Extended Data Figs. 8-16).

**Figure 3.**
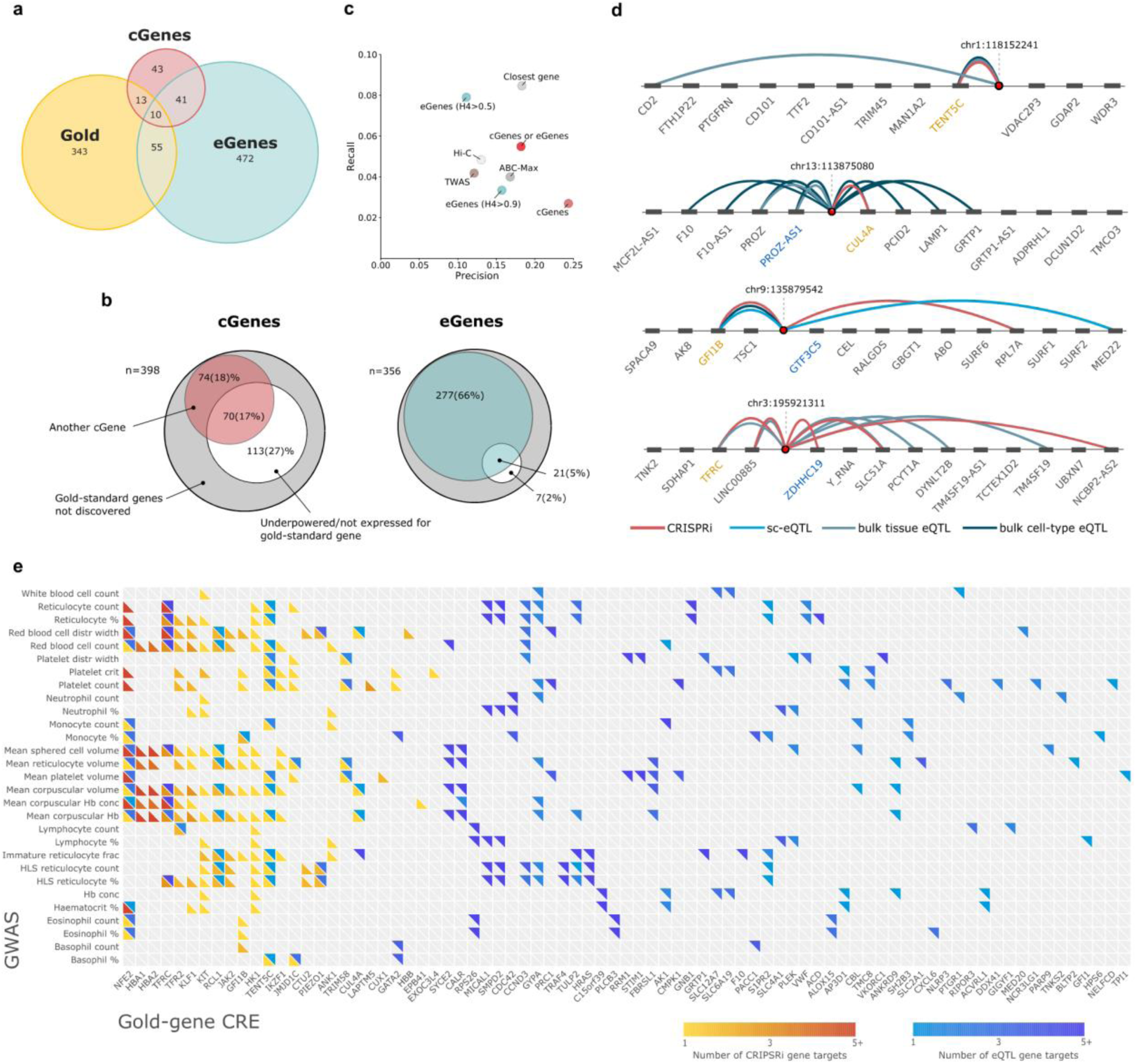
Discovery of gold-standard genes for blood traits. a, The overlap between cGenes, eGenes, and gold-standard genes**. b,** The gold-standard genes that were not identified as either cGenes or eGenes, with the proportion of undiscovered gold-standard genes for which eQTL and CRISPRi studies suggest a different causal gene at the same locus, as well as the proportion that remain undetected due to power limitations or lack of gene expression. **c,** Precision-recall for mapping GWAS variants to gold-standard causal genes (n = 1,075 CRE-gene pairs) within 1 Mb (n total = 15,699 CRE-gene pairs). Precision is the proportion of predicted positives that are true positives, while recall is the proportion of true positives that are correctly identified. **d**, Example loci showing the position of a GWAS *variant* and its gene associations as identified by different study types. Gold labels indicate the gold-standard gene in each locus, while blue labels mark the closest gene to the variant. If no blue label is present, the gold-standard gene is also the closest gene. **e,** Heatmap showing for each gold-standard gene intersecting a cGene and/or eGene (x-axis) and GWAS trait that the CRE mapping to that gene is associated to (y-axis), the number of genes located at the same locus identified by either CRISPRi perturbation screens (warm colours) or eQTL analyses (cool colours).

Next, we analyzed loci at which the gold-standard gene was not identified to understand why it was missed. In eQTL analysis, lack of power is an infrequent explanation, as only 7% of the loci had simulated statistical power <0.8. Again, other eGenes were detected in 84% of the loci, which may lead to a misleading inference. Conversely, in CRISPRi analysis, as much as 43% of gold-standard genes were either underpowered in CRISPRi study simulations (15%) or not expressed in K562 cells (29%), and in 37% of the loci, a different cGene was identified (Fig. 3b). We note that both eQTL and CRISPRi power calculations assume uniform effect sizes across CRE–gene pairs; however, distal or more constrained genes likely exhibit weaker regulatory effects, leading to overestimation of power for these targets. These results indicate distinct shortcomings in both approaches in the identification of causal genes for complex traits.

Assuming gold-standard genes represent true causal targets within a locus, we established a set of 1,075 causal CRE–gene pairs among 15,699 candidates located within ±1 Mb of each GWAS variant (Methods) to analyze the performance across target gene mapping strategies. cGenes achieved the highest precision (24%) but with the lowest recall (3%, Fig. 3c). eQTL colocalization identified more causal genes (recall = 8%) at lower precision (11%), and tightening the threshold to PP.H4 > 0.9 raised the precision to 16% while reducing recall to 3%. Assigning each GWAS variant to its nearest gene provided the greatest recall (9 %) with good precision (18%). Hi-C interactions and activity-by-contact (ABC) scores exhibited intermediate precision–recall trade-offs, with TWAS having a similar performance (Fig. 3c). Combining both cGenes and eGenes (PP.H4 > 0.9) offered increased recall (6%), while maintaining good precision (18%).The respective importance of precision or recall depends on the downstream applications. However, the largely non-overlapping subsets of gold-standard genes identified by the two methods point to their orthogonal nature in discovering candidate causal genes for GWAS loci.

## Discussion

We present the first systematic evaluation of the performance of the increasingly common CRISPRi approach for elucidating gene targets of GWAS loci, and contrast it to eQTL analysis. Our results revealed that the gene targets and GWAS CREs captured by each approach are often different. Some discordance is expected due to the differences in the designs of the analyzed studies, which rely on CRISPRi silencing in a cancer cell line versus genetic variation in primary cells. Furthermore, this work compares the results of the first generation of CRISPRi studies to results from more than a decade of eQTL work. The discoveries of the first eQTL studies were not too dissimilar from the patterns of CRISPRi results here: limited by discovery power, biased towards strong regulatory effects close to the promoter, detected in a single cell type ^25,26^. However, the major investments in both CRISPRi and eQTL studies to identify *cis* target genes for GWAS loci creates a critical need to understand the yield from each approach to pursue optimal study designs and practical applications, as well as make accurate biological inferences.

Our results show that while the eQTL data analyzed here was highly sensitive to detect proximal and distal cis-target genes of GWAS loci, important gaps remain. While sc-eQTL analysis has recently emerged as an attractive opportunity to capture genetic regulatory effects across diverse cellular contexts, further investments to increase power are needed for efficient GWAS target gene identification. eQTL analyses also struggle to detect regulatory effects of GWAS variants at lower allele frequencies and at low effect sizes. Many gold-standard GWAS genes being picked up by eQTLs with small effect sizes indicates continued opportunity for discovery in this realm. This is critical, as disease-relevant genes are more dosage-sensitive, constrained and thus harbor mostly minute regulatory effects and/or rare variants^9,19^. This, however, creates a double-edged sword: analyses that are sufficiently powered to detect small disease-relevant effects will readily capture many other genes as well, leading to loss of specificity, also demonstrated by our data. The real strength of eQTL studies is the study of primary cells and tissues, which inherently capture regulatory effects in a context that is closer to the in vivo state than model systems. Our results show how matching the analyzed cell types for the different blood cell traits contributes to the yield in target discovery, motivating ongoing investments in analysis genetic regulatory effects across disease-relevant cellular contexts.

The studied CRISPRi data sets detected a smaller number of target genes, mostly proximal to the GWAS locus, with statistical power limiting the discovery of weaker effects that predominate distal regulatory effects. This is an important gap in the ability to capture novel insights and unexpected GWAS target genes, as genes within tens of kilobases of a GWAS locus would be the first candidates based on gene annotation alone. However, our results show that CRISPRi target genes capture gold-standard GWAS genes with high specificity, and are better at detecting target genes that are more selectively constrained and tightly regulated. Ongoing expansion of CRISPR studies to accessible primary cell types and to variant-level editing will be critical to capture directly analogous disease biology and to understand to which extent this can be imputed from more scalable model systems and perturbations. Furthermore, CRISPR activation can complement CRISPRi by enhancing the power to detect lowly expressed genes. While we anticipate further discoveries as the CRISPR field grows and matures, the improvements in sensitivity, resolution, and diversity are likely to come with some loss of specificity.

Multiple gene targets of CREs across different cell types is a core feature of genome function ^27–29^, and accurate identification of which of the many target of a GWAS locus is the true driver of disease biology, and is likely not going to be achieved by cataloguing CRE targets alone. Rather, it will require further gene prioritization based on their functional properties. However, we note that while the identification of causal GWAS genes is a major research focus due to their role as potential drug targets^30^, this is not the only outcome or objective for eQTL and CRISPRi studies that also provide invaluable, generalizable information about the biology of gene regulatory mechanisms.

Our work is the first empirical demonstration of how both the eQTL and CRISPRi approaches have substantial limitations and orthogonal strengths. This highlights the importance of systematic evaluation of analytical approaches to understand their current yield and future opportunities. The complementarity of approaches that rely on population variation and experimental perturbations is an exciting opportunity for joint biological discovery that would be difficult to achieve with any approach alone, providing inspiration for building a diverse and comprehensive toolkit for GWAS interpretation.

## Methods

### Data

#### GWAS data

We utilized the genome-wide association study (GWAS) results for 29 different blood cell traits from 361,194 white British participants in the UK Biobank and trans-ethnic meta-analyses for 15 hematological traits in 746,667 participants from the BCX consortium ^31^. These traits included measures such as white blood cell count, red blood cell count, hemoglobin concentration, hematocrit percentage, and various others listed in data S1.

#### eQTL data

A total of 51 eQTL studies ^5,13–15,32^, focusing on blood and/or blood cell traits were analyzed (data S2). Summary statistics and log Bayes factors for most studies were obtained from the eQTL Catalogue ^13^. Summary statistics for GTEx ^5^, the Multi-Ancestry Analysis of Gene Expression (MAGE) study ^14^, the INTERVAL Study ^32^, and Meta LCL ^22^ were acquired from their respective study data sources listed in data S2. OneK1K association summary statistics were generated through re-analysis of the individual-level data (see Supplementary Methods).

#### CRISPRi perturbation data

CRISPRi data were obtained from Morris et al. ^6^, Gasperini et al. ^7^, and Gschwind et al ^16^. The Morris and Gasperini studies involved high-MOI CRISPRi screens targeting enhancers in K562 cells and were reanalyzed using SCEPTRE ^33^ with the default “union” setting, which groups gRNAs targeting the same element for association testing. Gschwind et al. compiled a harmonized gold-standard dataset of 10,411 element–gene pairs in K562 cells by reanalyzing CRISPR-based perturbation data from multiple studies. The largest number of CRE-gene targets within the Gschwind et al. study consisted of data from Schraivogel et al 2020., which utilized TAP-seq, and Nasser et al 2021., which utilized CRISPR FlowFISH. The different CRISPR datasets applied different candidate element selection strategies, leading to different characteristics, however, the vast majority of the data (>95% of CRE-gene targets) were data comprised from Morris et al. and Gasperini et al. which share a similar experimental design. To account for CREs that were targeted by multiple studies and the tiling approach utilized in flowFISH experiments we combined the targets of gRNA that were within 2kb of each other.

#### Capture Hi-C data data

We obtained capture Hi-C data in K562 cells from Zhigulev et al. 2024 ^34^. HiCapTools ^35^ was used to call interactions in all samples. A detailed description of how the interactions were called is reported elsewhere. Briefly, at least four pairs supporting each interaction were required. Three Bonferroni-adjusted *P*-value cutoffs (0.1, 0.01, 0.001) were used for filter interactions. For consistency with the eQTL and CRISPRi datasets, we used the 0.1 threshold. To identify GWAS target genes, we intersected fine-mapped credible sets with interacting regions using bedtools. All HiCap results were mapped to the GRCh37 genome assembly.

#### Determination of CRISPRi target genes

To link CRISPRi perturbations to GWAS variants, we obtained the SNP coordinates of the genomic regions targeted by gRNAs from Morris et al. For Gasperini et al., gRNA targeting coordinates were determined by aligning spacer sequences to the GRCh37 reference genome and identifying their positions within the targeted enhancers. Next, we intersected these CRISPRi-targeted regions with fine-mapped GWAS variants within ±2 kb to determine which credible set variants were perturbed by a particular CRISPRi experiment.

#### Gold-standard gene list

Summary statistics for pLoF burden tests on 29 blood cell traits were obtained from Genebass ^36^. Protein-coding genes with a SKAT p-value < 5 × 10^-7^ and located within 1 Mb of GWAS CREs targeted by CRISPRi gRNAs were selected for analysis. In addition to these burden test genes, curated gene lists associated with rare inherited Mendelian blood disorders ^37,38^ that were also in cis (within 1 Mb) of a GWAS CRE were also included. Fisher’s exact tests were performed to assess the enrichment of overlap between Mendelian genes, burden test genes, cGenes, eGenes, and Hi-C gene targets. Hi-C, genes were filtered to protein-coding genes that were associated with a CRE within 1 Mb of a gold-standard gene.

#### Activity-by-contact gene predictions

Activity-by-Contact (ABC) enhancer–gene predictions were overlapped with the genomic coordinates of our CRISPR-targeted candidate regulatory elements (CREs) to identify putative target genes for each guide RNA. We first downloaded the ABC prediction file (AllPredictions.AvgHiC.ABC0.015.minus150.ForABCPaperV3.txt.gz; Nasser et al., 2021) from the Broad Institute FTP server. A subset of immune-relevant and hematopoietic cell types was specified, including K562 (Roadmap), B cells (ENCODE and BJAB), CD4⁺ and CD8⁺ αβ T cells (ENCODE and Corces 2016), CD14⁺ monocytes under baseline and stimulated conditions (ENCODE and Novakovic 2016), LPS-treated dendritic cells (Garber 2017), megakaryocyte–erythroid progenitors (Corces 2016), NK cells (Roadmap and Corces 2016), GM12878 (Roadmap), Jurkat (Engreitz), U937 (Engreitz), THP-1 monocytes/macrophages (Van Bortle 2017), and OCI-LY7 (ENCODE). The CRE-guide coordinate file was intersected with each cell-type-specific ABC file using bedtools intersect resulting in 466 gRNAs that intersected a ABC enhancer. Each gRNA coordinate was assigned to the gene with the max ABC score (ABC-Max) across all the cell types specified.

#### TWAS gene predictions

To identify the most significant transcriptome-wide association study (TWAS) gene in cis for each GWAS CRE test, we used results from the Rowland et al. who performed a TWAS for 29 blood-cell traits from the UK biobank ^39^. We retained only protein-coding genes and for each GWAS CRE, we defined a cis window as ±1 Mb around the gRNA’s genomic position. Within this window, we extracted all TWAS associations whose start or end coordinates overlapped the interval on the matching chromosome. If one or more associations were found, we retained the gene with the most significant association (i.e., the highest log10 p-value). TWAS associations were only retained if they passed a marginal significance threshold (as defined in the TWAS dataset). The resulting dataset consisted of the most significant cis TWAS gene for each CRISPR perturbation, and was used to evaluate the overlap between TWAS predictions and CRISPR-based regulatory inferences.

## Statistical analysis

### Statistical fine-mapping of UK Biobank blood cell traits

For each of the 29 UK Biobank blood trait GWASs, statistical fine-mapping was conducted using Sum of Single Effects regression (SuSiE), as implemented in the susieR R package ^40^. We defined 899 disjoint fine-mapping regions (Supplementary Methods) containing genome-wide significant variants (*P* < 6.6 × 10^-9^). We generated linkage disequilibrium (LD) matrices from a subset of 50,000 UK Biobank white British participants (UK Biobank accession code: 47976) using PLINK. We extracted all variants in the 95% credible set with a posterior inclusion probability (PIP) > 0.01.

### Colocalization with UKBB GWAS variants

We performed colocalization between 29 UK Biobank blood trait GWASs and 49 eQTL studies of European ancestry. Pairwise colocalization was conducted using SuSiE COLOC ^41^ between each blood trait GWAS and eQTL study. When available, we used fine-mapped eQTL data from the eQTL Catalogue; otherwise, eQTL data were fine-mapped before colocalization using in-sample LD matrices. To ensure consistency in genomic positions, all data were mapped to the GRC37 using bcftools liftover ^42^. Colocalization was performed within regions meeting the thresholds of p < 6.6 × 10⁻⁹ for GWAS associations and p < 1 × 10^-5^ for eQTL signals. SuSiE COLOC was conducted with default settings, and if convergence failed at a particular locus, colocalization was performed using COLOC ^43^ under the assumption of a single causal variant. Additionally, we conducted colocalization between 15 blood cell traits from the BCX GWAS ^31^ consortium and the multi-ethnic eQTL study MAGE ^14^. For this analysis, COLOC was performed under the single causal variant assumption without requiring additional LD information.

### Statistical testing

To compare the number of enhancers between gene sets, we used Poisson regression. Differences in Episcore, pHaplo, gene expression levels in K562 and GTEx whole blood, and MAF were assessed with Gaussian regression, while differences in pLI were assessed with binomial regression. All gene-level models were adjusted for gene length.

We applied the Wilcoxon rank-sum test to compare eQTL colocalization probabilities and Fisher’s exact test to evaluate the number of genes with estimated CRISPRi or eQTL power greater than 0.8. Chi-squared tests were used to assess differences in categorical distributions, including the number of genes per locus and gene–CRE distances. All statistical tests were two-sided. Associations between log-fold change and TSS distance were tested using linear regression, adjusting for the number of cells bearing each gRNA.

### CRISPRi power calculation

We estimated the power to detect reductions in gene expression for each of the perturbed CRE–target gene pairs using a modified version of the approach published by Gschwind et al. 2023^16^. For the Morris et al. and Gasperini et al. datasets, we first applied the estimateDispersions function (DESeq2, v1.46.2) to estimate mean and overdispersion parameters across cells for each gene. Using these estimates, we sampled expression values from a negative binomial distribution, introducing a fixed reduction in expression (15%) for each perturbed gene linked to a given CRE. We calculated power for effect sizes 10%, 15%, 20%, 25%, 30%, 40%, 50% reduction in expression. An effect size of 25% was determined based on the number of significant empirical CRISPRi CRE-gene associations observed within the data, in comparison to the number significant empirical eQTL variant gene associations. We sampled the same number of cells for each perturbation as the original datasets and aggregated them to ensure that perturbed and non-perturbed cell counts remained consistent. We next ran SCEPTRE in high-MOI mode—without any extra quality filtering—to assess each CRE–gene pair for a significant decrease in expression, correcting P-values across all pairs within each dataset with the Benjamini–Hochberg procedure. The entire simulation was repeated 10 times, and a CRE–gene pair was considered powered for detection if a significant expression change was observed at FDR < 10% in at least 80% of simulations. For the remaining datasets, power estimates produced by Gschwind et al. 2023 were used ^16^.

### Estimation of equivalent eQTL effect size for CRISPRi perturbations

Empirical power analyses indicated that approximately 80% of significantly perturbed CREs exhibited ≥80% statistical power at an estimated 25% reduction in gene expression relative to non-targeting control (NTC) cells. To translate this level of repression into an equivalent eQTL effect size, we used allelic fold-change (aFC) estimates from GTEx v10 Whole Blood eQTLs. eGenes were restricted to those with q-values ≤0.05. We identified variants whose rounded aFC values approximated log2(0.75), corresponding to a 25% decrease in expression, while excluding variants whose 95% confidence intervals encompassed zero. The mean β-coefficient among these variants was 0.27, which we used as the eQTL effect size comparable to the CRISPRi perturbation magnitude.

### eQTL power calculation

For genes with a mean count greater than 5, power is estimated analytically using an F-test. Power in this context depends on sample size n, the significance threshold α (*P* = 1 × 10^-3^), and the proportion of variance explained (*q²*) by the variant in the population. Assuming genotypes are in Hardy-Weinberg equilibrium, *q²* is given by:

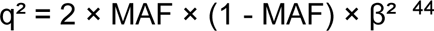

The minor allele frequency (MAF) refers to the GWAS variant, with β set to 0.25 and the phenotype (gene expression) expressed in standard deviation units.. For lowly expressed genes, power is determined through simulations using scPower ^45^. In sc-eQTL studies, the overall power is the product of eQTL power and expression probability, which quantifies the l likelihood of a gene being detected in a given cell type ^45^. Expression probability is determined by sample size, the number of cells per cell type, cell type frequencies, and the proportion of individuals in which a gene is expressed. Read depth is implicitly accounted for through UMI counts. When comparing power across studies and methods, we used 13,874 protein-coding genes that passed QC in the INTERVAL study as the reference set, assumed to be expressed across all studies. For a detailed description of power calculations, refer to the Supplementary Methods.

### Trans-eQTL calling

To identify GWAS CREs with trans-eQTLs, we selected *cis*-eQTL variants for genes that displayed significant trans-effects in CRISPRi studies and were cis-targets of GWAS CREs. Trans-eQTL analysis was then performed for these cis-eQTL variants across Azimuth level 1 predicted cell types in the OneK1K dataset using tensorQTL ^46^. Trans associations within a ±5 Mb window that met a significance threshold of FDR < 0.05 were retained. Additionally, trans-eQTL results from a meta-analysis of Lymphoblastoid Cell Lines (LCLs) in MetaLCL ^22^ were incorporated. Using the same approach, we identified cis-eQTL variants for genes with CRISPR-derived trans networks that were also GWAS targets based on MAGE data ^14^. We then extracted trans associations for these cis-eQTLs from the MetaLCL results and retained those with FDR < 0.05 to further assess their trans-regulatory effects.

### Trans-eQTL comparison

To assess concordance of trans-effects between methods, we first identified significant trans-cGene associations (FDR < 0.10) from Morris et al. and Gasperini et al. using SCEPTRE, filtered to cis-genes linked to GWAS CREs. Trans-eGene effect sizes were then aligned to the allele associated with decreased expression of the cis-gene. We then tested the correlation of effect sizes across overlapping trans-genes using Pearson correlation, adjusting p-values for multiple testing with the Benjamini-Hochberg procedure. Enrichment analyses were conducted using a permutation-based Fisher’s exact test to estimate odds ratios and confidence intervals, with multiple testing correction applied similarly.

### Gene properties

Enhancer and super-enhancer counts per gene were derived from GeneHancer v5 ^47^ ^47^, with filtering applied to retain enhancers with a score ≥ 0.7 and gene associations with a “link_score” ≥ 7. Haploinsufficiency (pHaplo) scores and Episcore values were obtained from Collins et al ^48^. pLI scores were retrieved from gnomAD v3, and gene expression levels (TPM) for whole blood were obtained from GTEx version 8. Hi-C data for enhancer-promoter interactions in K562 was obtained from Zhigulev et al ^34^.

### Calculation of Precision and Recall for gene mapping approaches

To assess the sensitivity and specificity of CRISPRi and eQTLs in identifying the correct target gene, we evaluated all genes within ±1 Mb of each GWAS CRE that contained a gold-standard gene and at least one significant cGene or eGene. Any gold-standard gene at each locus was treated as the true causal gene. 497 GWAS CREs contained at least one gold-standard gene within 1 Mbp, resulting in 1,075 causal CRE-gene connections were determined out of a total of 13,000 CRE-gene links. We limited our analysis to only protein coding genes as the majority of the gold-standard genes were protein coding. A CRE–gene match to the gold-standard was counted as a true positive (TP); identification of a different gene as a false positive (FP); failure to identify the gold-standard as a false negative (FN); and all correctly excluded non-gold genes as true negatives (TN). Recall was calculated as TP / (TP + FN), and precision as TP / (TP + FP).

## Acknowledgments

This research has been conducted using the UK Biobank Resource under Application Number 47976. The authors would like to thank Angli Xue for valuable discussions and for providing the OneK1K raw VCF file and cell-type classifications.

This work was supported by funding from the National Institutes of Health R01MH106842, 24HG012090, R01HL168247 and R01HG012790, the European Research Council (ERC) under the European Union’s Horizon 2020 research and innovation programme (Grant agreement no. 101043238), and Knut and Alice Wallenberg Foundation grant KAW 2023.0337. A part of the computations were enabled by resources in project (SNIC 2022-5-560 and NAISS 2023-5-517) provided by the National Academic Infrastructure for Supercomputing in Sweden (NAISS) at UPPMAX, funded by the Swedish Research Council through grant agreement no. 721 2022-06725.

## Author contributions

S.G., J.A.M., and T.L. conceived and designed the project. S.G. performed most of the data analysis. J.P. and S.G. performed power analyses and advised on analysis methods. W.O. recalled eQTLs from OneK1K. J.A.M., S.G., and T.L. provided feedback on the study design and data analysis. J.A.M. and T.L. supervised the work. S.G. and T.L. wrote the manuscript with contributions from other authors. All authors read, provided feedback, and approved the manuscript.

## Data availability

GWAS CRE and eGene and cGene target results are available within the supplementary tables of this manuscript. The fine-mapped credible sets and log-Bayes factors for the UKBB, GTEx, and OneK1K will be available for download upon publication at: https://drive.google.com/drive/folders/1UdnlUs-xtutWk_hSofk1iX0tnS64P9dO?usp=drive_link.

The summary statistics of the recalled eQTLs of the OneK1K dataset will also be available upon publication at: https://drive.google.com/drive/folders/1Q3Zz4EFvvgn4gYllBJQl-q8605AYOhSO?usp=drive_link.

Reanalyzed CRISPRi data URLs can be found within data S3. We obtained summary statistics and log-Bayes factors for eQTL studies from the eQTL catalog ^13^. GWAS summary statistics for 44 blood cell traits were downloaded from the Neale Lab (http://www.nealelab.is/uk-biobank/) and Lettre Lab (http://www.mhi-humangenetics.org/en/resources/). The datasets used in this study are publicly available from various sources. GeneHancer v5.10 provides enhancer size and the number of enhancers per gene, this dataset is available upon request from GeneCards: https://www.genecards.org/Download/File?file=GeneHancer_v5.10.gff.

GTEx v8 transcript-level TPM data was obtained from GTEx Analysis v8 and can be accessed at https://storage.googleapis.com/gtex_analysis_v8/rna_seq_data/GTEx_Analysis_2017-06-05_v8_RSEMv1.3.0_transcript_tpm.gct.gz. The pHaplo and pTriplo scores, which estimate the probability of a gene being haploinsufficient or triplosensitive, along with Episcore values, were obtained from https://ars.els-cdn.com/content/image/1-s2.0-S0092867422007887-mmc7.xlsx.

The pLI and LOEUF scores were retrieved from gnomAD v3 and are available at https://gnomad.broadinstitute.org/. Hi-C data for enhancer-promoter interactions in blood cell lines, including K562, was obtained from https://kth-my.sharepoint.com/:f:/g/personal/pelinak_ug_kth_se/Eq0b5SU6LGVFj4My7hiQiwUB5gmQuHEL-unZRrBCRa19ig?e=MinXnm.

## Code availability

CRISPRi power calculations are available at: https://github.com/jasperpanten/sc_crispr_simulations

All other analysis codes are available at: https://github.com/PandaPowell/CRISRPi-eQTLs-Manuscript

