## Supplementary Figures for "CRISPRi perturbation screens and eQTLs provide complementary and distinct insights into GWAS target genes"

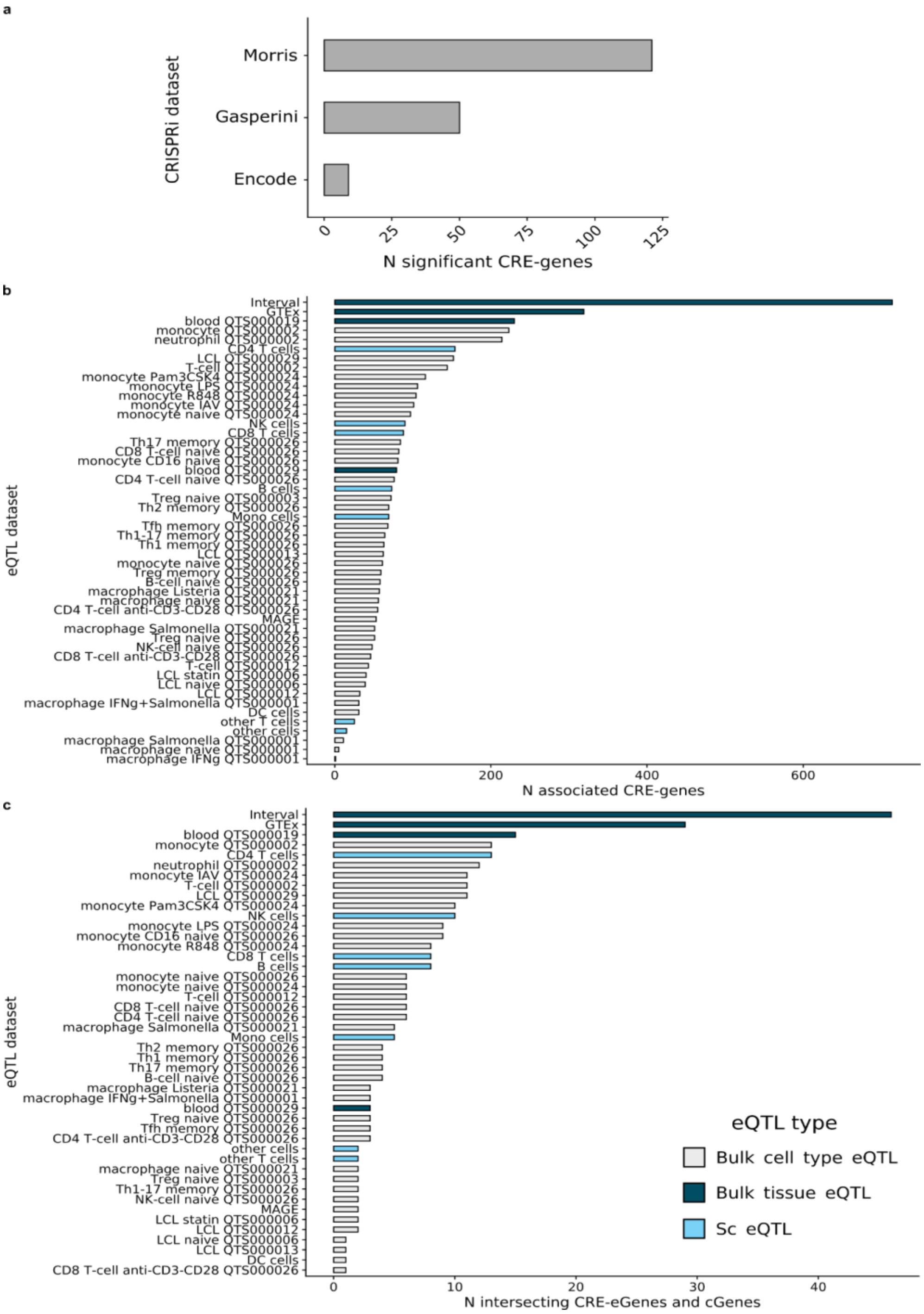


***Fig S1 - Gene discovery by eQTL and CRISPRi studies. a,*** *The number of significant CRE-genes per CRISPRi dataset.* ***b****, The number of variant gene associations by eQTL study, codes represent eQTL catalogue study IDs.* ***c****, The number of intersecting eGenes and cGenes by eQTL study.*

a

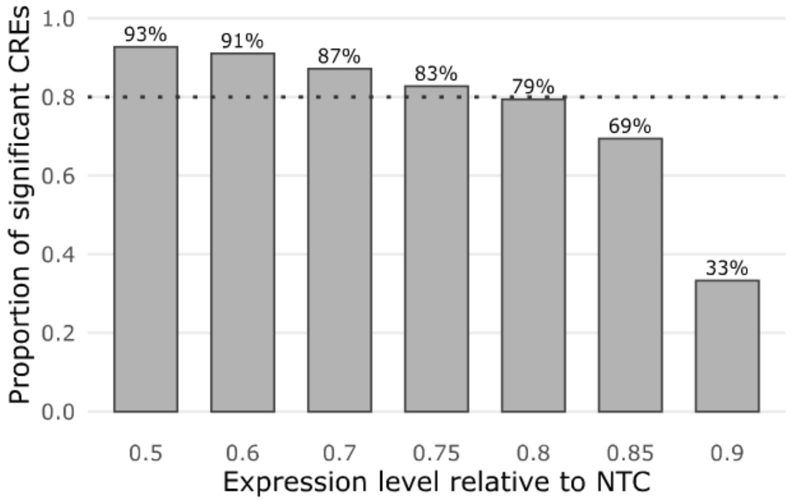


b

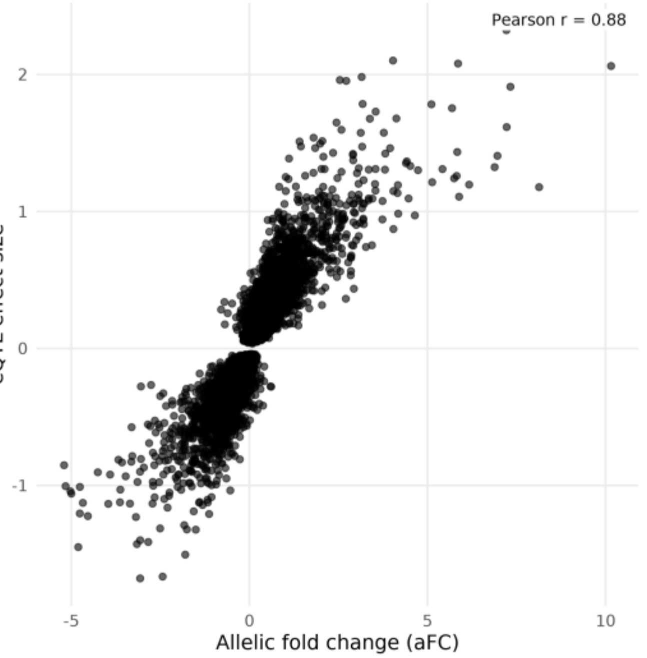


c

| Target aFC | Target % change | Mean eQTL slope (β) |
|---|---|---|
| -1.000 | -50.0% | -0.381 |
| -0.515 | -30.0% | -0.282 |
| -0.415 | -25.0% | -0.265 |
| -0.322 | -20.0% | -0.249 |
| -0.234 | -15.0% | -0.235 |
| -0.152 | -10.0% | -0.223 |

***Fig S2 – CRISPRi and eQTL power effect size selection**. **a**, Proportion of significant CREs across CRISPRi effect sizes. Bars show the fraction of CRE–gene pairs reaching significance at each relative expression level (CRISPRi vs. control). The dotted line marks the 80% power threshold. **b**, Allelic fold-changes (aFC) and eQTL effect sizes (linear regression beta) for significant GTEx v10 Whole Blood eGenes. **c,** fixed target expression changes (−10 % to −50 %) used in downstream simulations, together with their aFC and eQTL effect size equivalents within for GTEx v10 Whole Blood eGenes.*

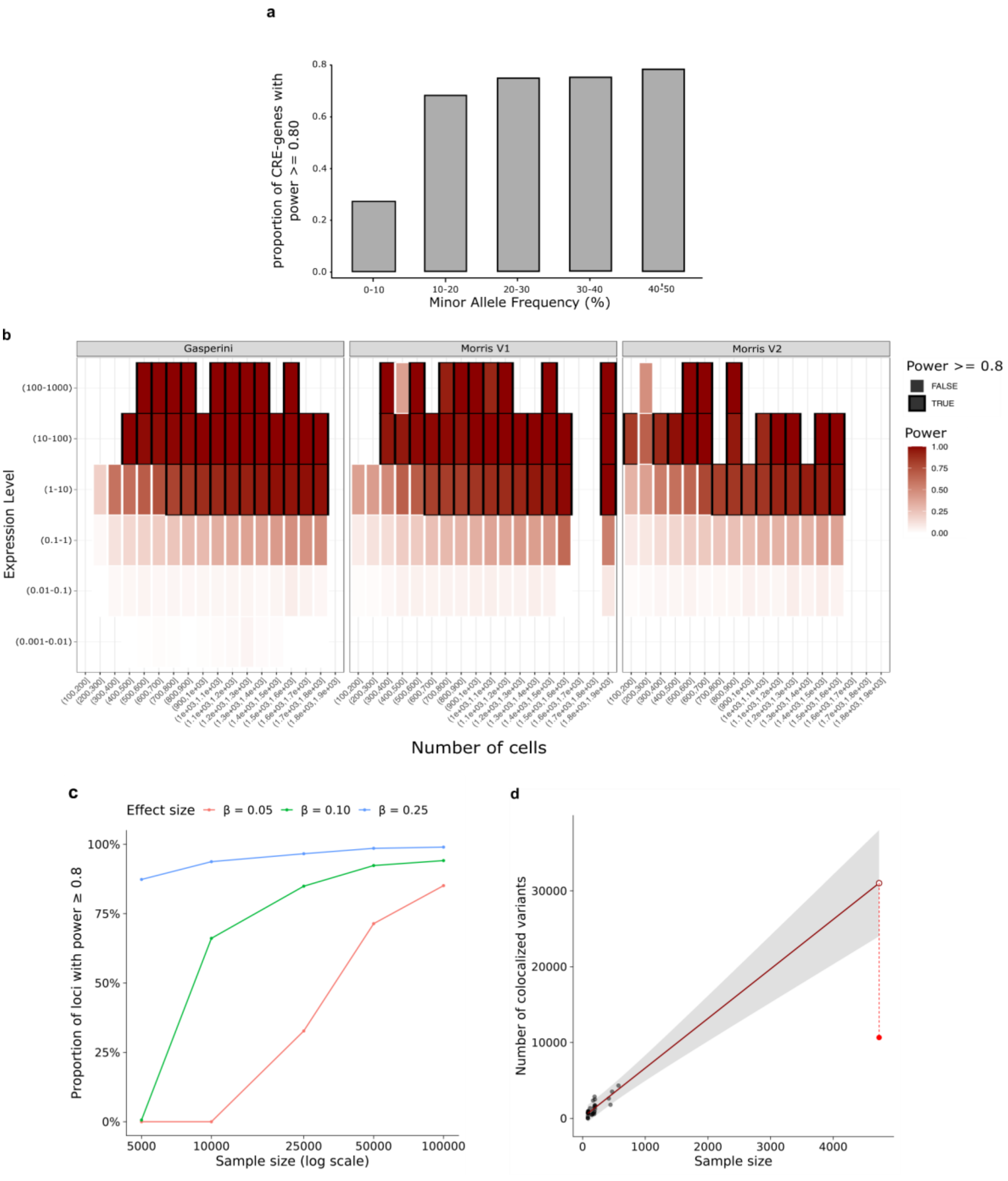


***Fig S3 - Sample size and power of eQTL and CRISPRi studies. a****, Estimated detection power for eQTLs across minor allele frequencies.* ***b,*** *Estimated CRISPRi power as a function of the number of cells and gene expression level. Morris data consisted of two experimental stages, version 1 and 2.* ***c****, The*

*proportion of eQTL loci achieving ≥80% power is shown as a function of sample size for effect sizes (β) of 0.05, 0.10, and 0.25. Curves represent mean power estimates derived from simulations across loci, illustrating how larger sample sizes are required to achieve adequate power for smaller effect sizes.* **d,** *Relationship between study sample size and the empirical number of colocalized loci. Points show individual studies; the largest cohort (INTERVAL) is highlighted in red. The solid line is the leave-one-out linear fit (ordinary least squares) trained on all cohorts except the largest; the shaded band is its 95% prediction interval. The open circle at the maximal sample size marks the model-predicted value; the dashed segment shows the residual to the observed value. The largest cohort lies outside the linear model's 95% prediction band and is influential in the full linear fit (high studentized residual (−5.39; $P = 3.45 \times 10^{-6}$) and Cook's distance (D=338).*

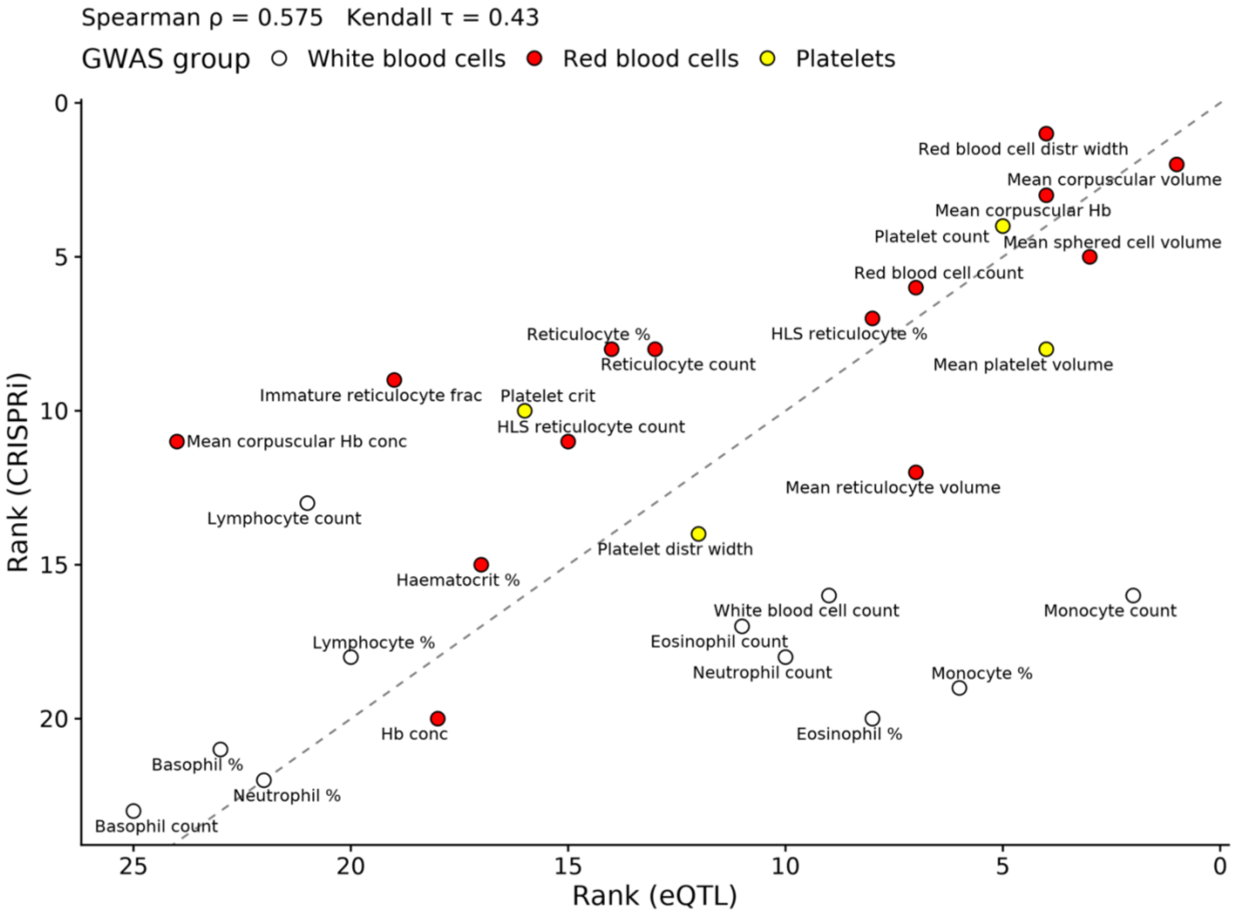


***Fig S4 – Rank concordance of trait associations.*** *Each point represents a GWAS trait, ranked independently by the number of significant CRE–gene associations identified in CRISPRi and eQTL analyses. Points along the diagonal (dashed line) indicate traits with consistent relative rankings between methods, whereas deviations reflect traits more strongly represented in one approach than the other. Axes are reversed so that higher ranks (rank 1) appear in the top-right corner. Colors denote the corresponding GWAS trait groups (e.g., red blood cells, white blood cells, platelets). Spearman's ρ and Kendall's τ measure the strength of rank concordance between the two datasets.*

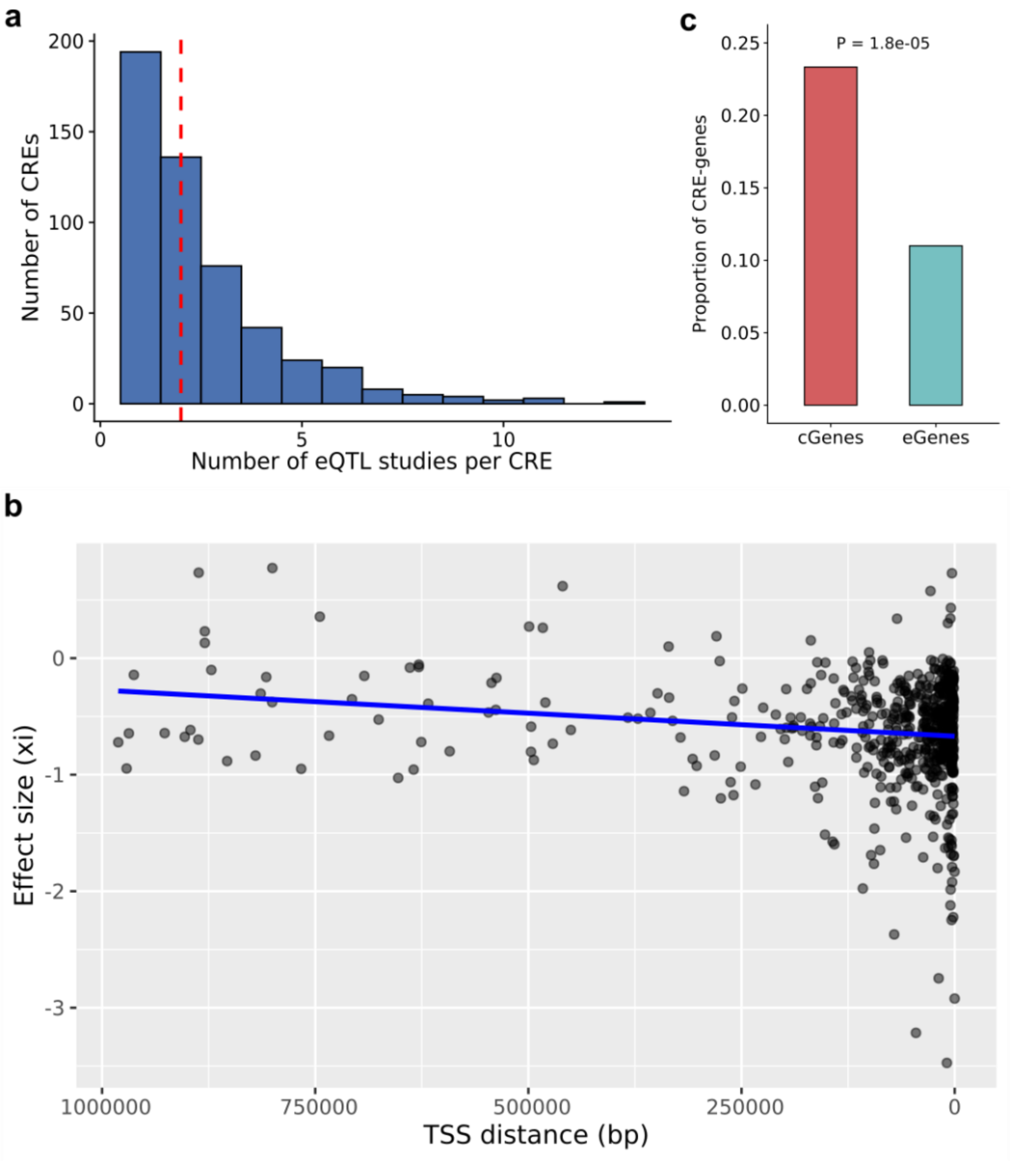


***Fig S5 - Comparative power and genomic features influencing cGene–eGene overlap. a****, The number of eQTL studies per CRE that map that CRE to a unique gene.* ***b****, Log-fold change decreases with CRE–TSS absolute distance across Morris and Gasperini datasets. A regression line (blue) adjusts for the number of cells per gRNA.* ***c****, Proportion of Hi-C target genes that intersect with gold-standard genes.*

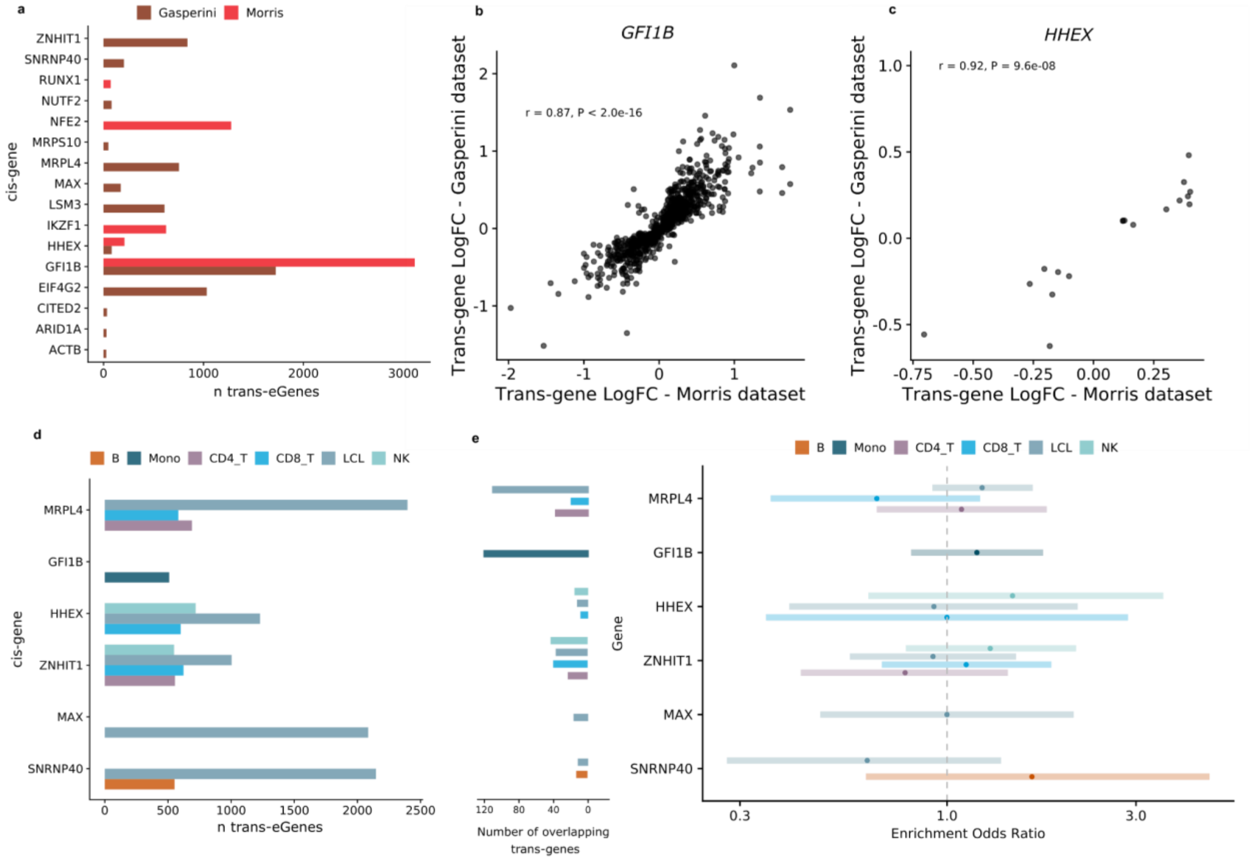


***Fig S6 - Trans-genes analysis of GWAS CREs a,*** *Number of significant trans-cGenes per cis-cGene.* ***b****, Correlation of GFI1B-targeting trans-cGene effects across Morris and Gasperini CRISPRi screens.* ***c****, Correlation of HHEX-targeting trans-cGene effects across Morris and Gasperini CRISPRi screens.* ***d****, Number of significant trans-eGenes per cis-eGene.* ***e****, Trans-gene overlap analysis, the y-axis indicating the cis-genes through which trans-effects occur. The left-hand side of the plot shows the number of overlapping trans eGenes and cGenes, and the enrichment odds ratio and its confidence interval of the effect sizes are shown on the right-hand side.*

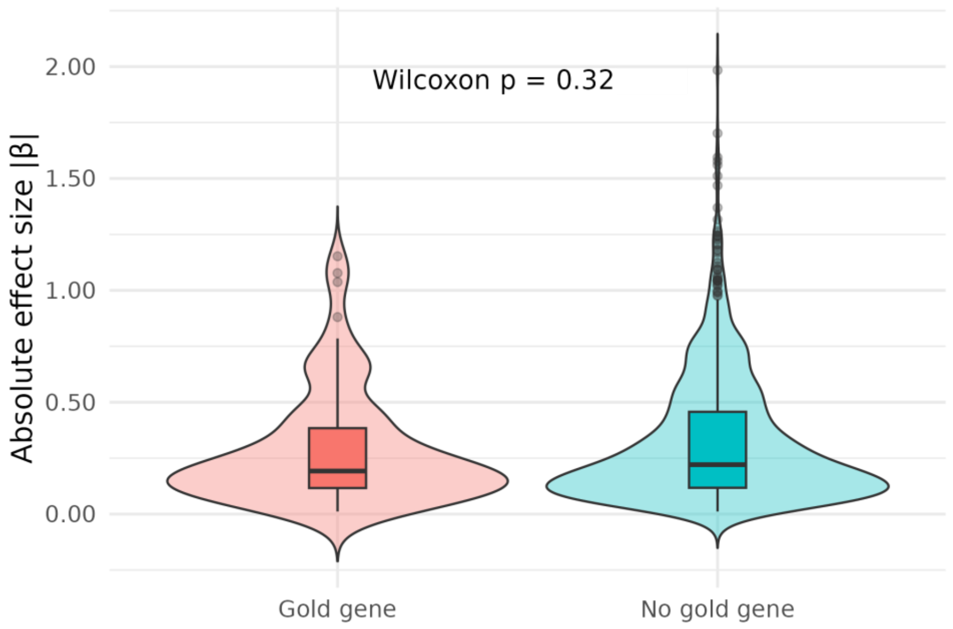


***Fig S7 - Distribution of eQTL effect sizes for gold-standard and non–gold-standard genes -*** *Violin plots show the distribution of absolute eQTL effect sizes β for variants intersecting gold-standard genes (left) and those not overlapping gold-standard genes (right). Median and interquartile ranges are indicated by boxplots.*

***Fig S8 – 16 - Locus heatmaps of cGenes and eGenes intersecting gold-standard genes.*** *Each panel shows a single GWAS CRE locus. Rows correspond to individual eQTL or CRISPRi studies; columns list all genes in the locus ranked by distance to the CRE. Cell colour denotes the –$log_{10}$ P value (red = stronger association). CRE position label marks the position of the variant in relation to the genes within the locus. Gold text marks the gold-standard gene, blue text marks the gene closest to the CRE (gold only, if the two coincide).*

**8.**

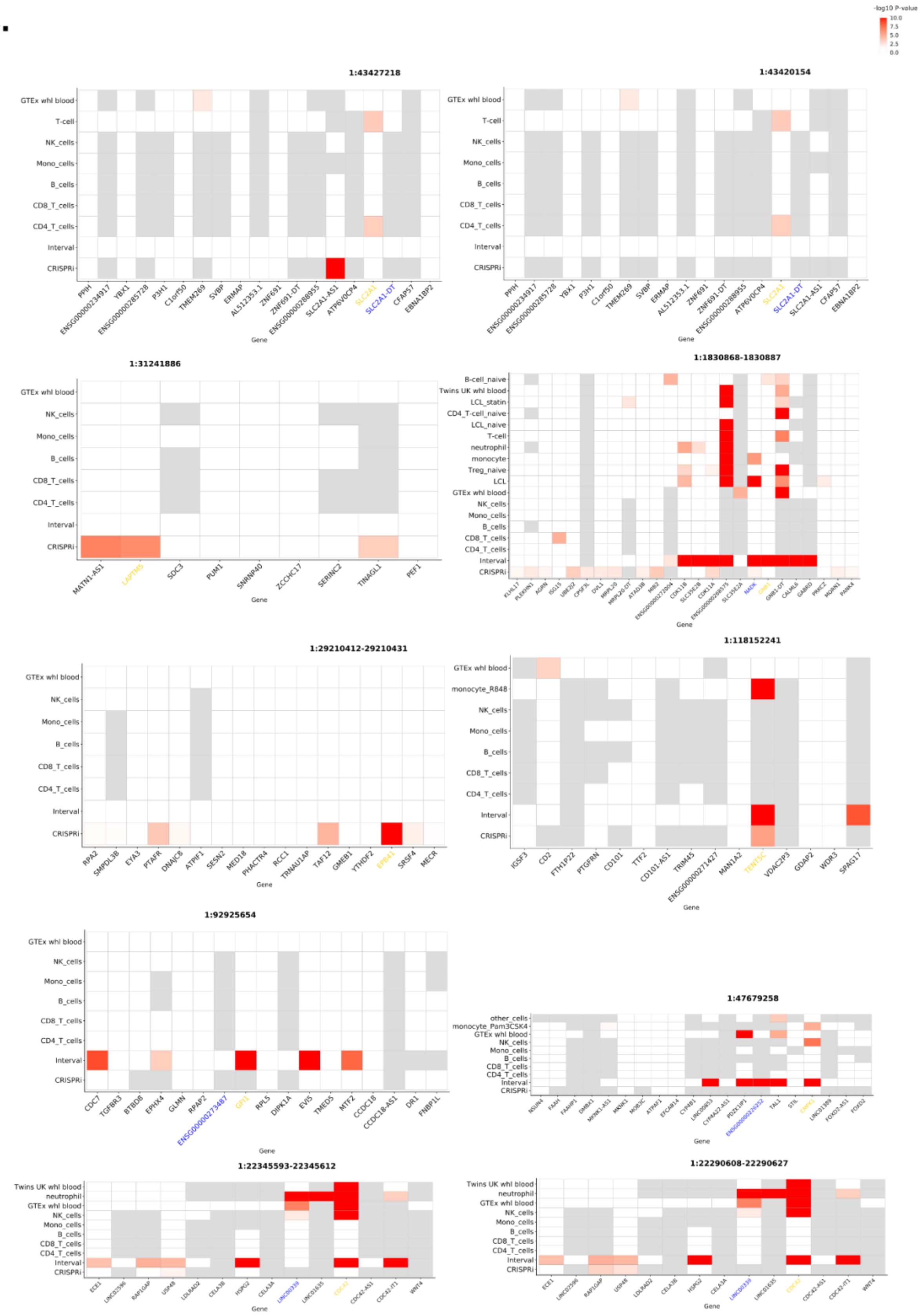
-log10 P-value
10.0
7.5
5.0
2.5
0.0
1:43427218
GTEx whl blood
T-cell
NK_cells
Mono_cells
B_cells
CD8_T_cells
CD4_T_cells
Interval
CRISPRi
PPIH
ENSG00000234917
YBX1
ENSG00000285728
P3H1
C1orf50
TMEM269
SVBP
ERMAP
AL512353.1
ZNF691
ZNF691-DT
ENSG00000288955
SLC2A1-AS1
ATP6V0CP4
SLC2A1
SLC2A1-DT
CFAP57
EBNA1BP2
Gene
1:43420154
1:31241886
MATN1-AS1
LAPTM5
SDC3
PUM1
SNRNP40
ZCCHC17
SERINC2
TINAGL1
PEF1
1:1830868-1830887
B-cell_naive
Twins UK whl blood
LCL_statin
CD4_T-cell_naive
LCL_naive
neutrophil
monocyte
Treg_naive
LCL
1:29210412-29210431
RPA2
SMPDL3B
EYA3
PTAFR
DNAJC8
ATPIF1
SESN2
MED18
PHACTR4
RCC1
TRNAU1AP
TAF12
GMEB1
YTHDF2
EPB41
SRSF4
MECR
1:118152241
monocyte_R848
IGSF3
CD2
PTGFRN
FTH1P22
CD101
TTF2
CD101-AS1
TRIM45
ENSG00000271427
MAN1A2
TENT5C
VDAC2P3
GDAP2
WDR3
SPAG17
1:92925654
CDC7
TGFBR3
BTBD8
EPHX4
GLMN
RPAP2
ENSG00000273487
GFI1
RPL5
DIPK1A
EVI5
TMED5
MTF2
CCDC18
CCDC18-AS1
DR1
FNBP1L
1:47679258
other_cells
monocyte_Pam3CSK4
1:22345593-22345612
1:22290608-22290627

**9.**

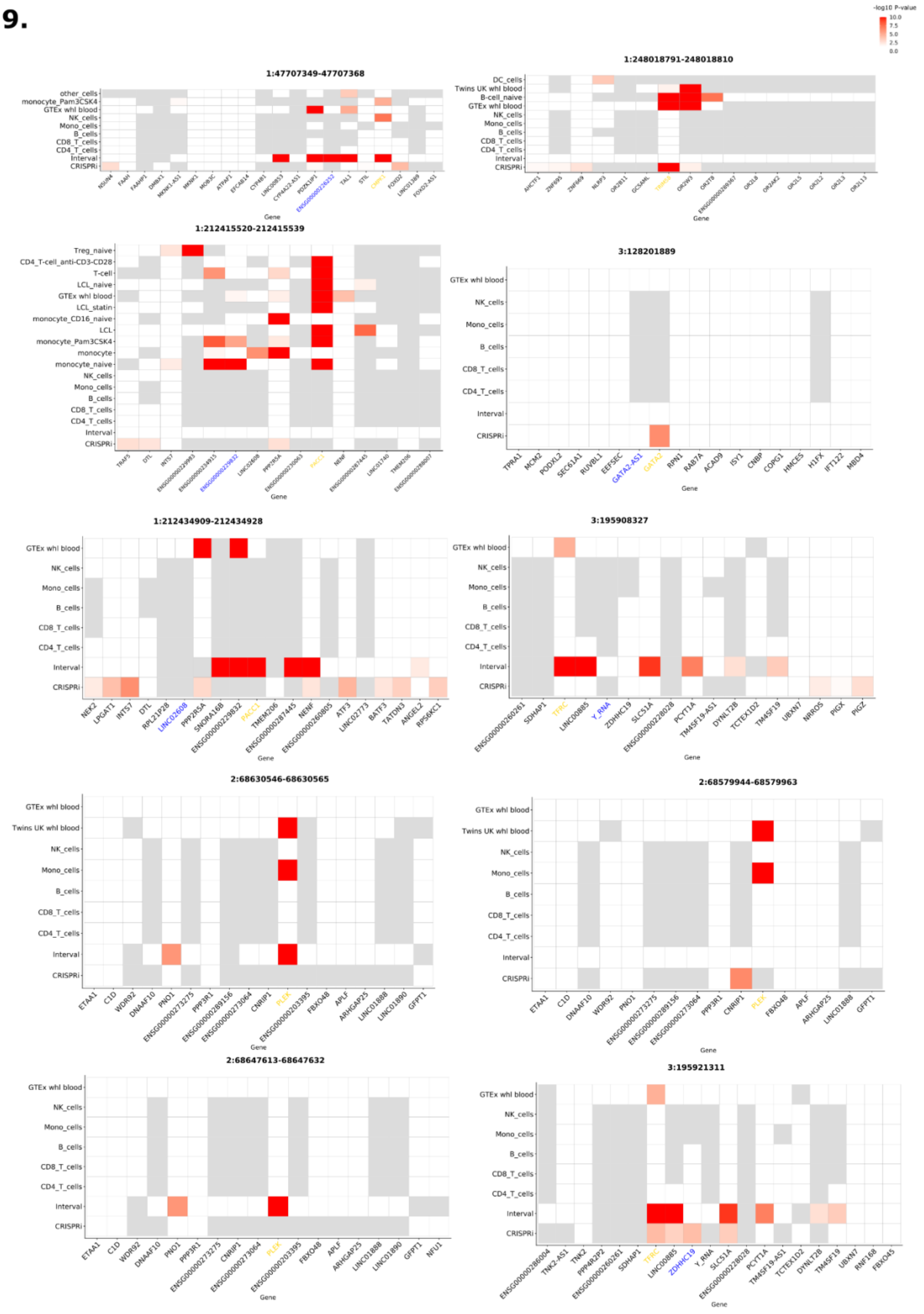
-log10 P-value
10.0
7.5
5.0
2.5
0.0
1:47707349-47707368
other_cells
monocyte_Pam3CSK4
GTEx whl blood
NK_cells
Mono_cells
B_cells
CD8_T_cells
CD4_T_cells
Interval
CRISPRi
Gene
1:248018791-248018810
DC_cells
Twins UK whl blood
B-cell_naive
GTEx whl blood
NK_cells
Mono_cells
B_cells
CD8_T_cells
CD4_T_cells
Interval
CRISPRi
Gene
1:212415520-212415539
Treg_naive
CD4_T-cell_anti-CD3-CD28
T-cell
LCL_naive
GTEx whl blood
LCL_statin
monocyte_CD16_naive
LCL
monocyte_Pam3CSK4
monocyte
monocyte_naive
NK_cells
Mono_cells
B_cells
CD8_T_cells
CD4_T_cells
Interval
CRISPRi
Gene
3:128201889
GTEx whl blood
NK_cells
Mono_cells
B_cells
CD8_T_cells
CD4_T_cells
Interval
CRISPRi
TPRA1
MCM2
PODXL2
SEC61A1
RUVBL1
EEFSEC
GATA2-AS1
GATA2
RPN1
RAB7A
ACAD9
ISY1
CNBP
COPG1
HMCES
H1FX
IFT122
MBD4
Gene
1:212434909-212434928
GTEx whl blood
NK_cells
Mono_cells
B_cells
CD8_T_cells
CD4_T_cells
Interval
CRISPRi
NEK2
LPGAT1
INTS7
DTL
RPL21P28
LINC02608
PPP2R5A
SNORA16B
ENSG00000229832
PACC1
TMEM206
ENSG00000287445
NENF
ENSG00000260805
ATF3
LINC02773
BATF3
TATDN3
ANGEL2
RPS6KC1
Gene
3:195908327
GTEx whl blood
NK_cells
Mono_cells
B_cells
CD8_T_cells
CD4_T_cells
Interval
CRISPRi
ENSG00000260261
SDHAP1
TFRC
LINC00885
Y_RNA
ZDHHC19
SLC51A
ENSG00000228028
PCYT1A
TM4SF19-AS1
DYNLT2B
TCTEX1D2
TM4SF19
UBXN7
NRROS
PIGX
PIGZ
Gene
2:68630546-68630565
GTEx whl blood
Twins UK whl blood
NK_cells
Mono_cells
B_cells
CD8_T_cells
CD4_T_cells
Interval
CRISPRi
ETAA1
C1D
WDR92
DNAAF10
PNO1
ENSG00000273275
PPP3R1
ENSG00000289156
ENSG00000273064
CNRIP1
PLEK
ENSG00000203395
FBXO48
APLF
ARHGAP25
LINC01888
LINC01890
GFPT1
Gene
2:68579944-68579963
GTEx whl blood
Twins UK whl blood
NK_cells
Mono_cells
B_cells
CD8_T_cells
CD4_T_cells
Interval
CRISPRi
ETAA1
C1D
DNAAF10
WDR92
PNO1
ENSG00000273275
ENSG00000289156
ENSG00000273064
PPP3R1
CNRIP1
PLEK
FBXO48
APLF
ARHGAP25
LINC01888
GFPT1
Gene
2:68647613-68647632
GTEx whl blood
NK_cells
Mono_cells
B_cells
CD8_T_cells
CD4_T_cells
Interval
CRISPRi
ETAA1
C1D
WDR92
DNAAF10
PNO1
PPP3R1
ENSG00000273275
CNRIP1
ENSG00000273064
PLEK
ENSG00000203395
FBXO48
APLF
ARHGAP25
LINC01888
LINC01890
GFPT1
NFU1
Gene
3:195921311
GTEx whl blood
NK_cells
Mono_cells
B_cells
CD8_T_cells
CD4_T_cells
Interval
CRISPRi
ENSG00000286004
TNK2-AS1
TNK2
PPP4R2P2
ENSG00000260261
SDHAP1
TFRC
LINC00885
ZDHHC19
Y_RNA
SLC51A
ENSG00000228028
PCYT1A
TM4SF19-AS1
TCTEX1D2
DYNLT2B
TM4SF19
UBXN7
RNF168
FBXO45
Gene

10.

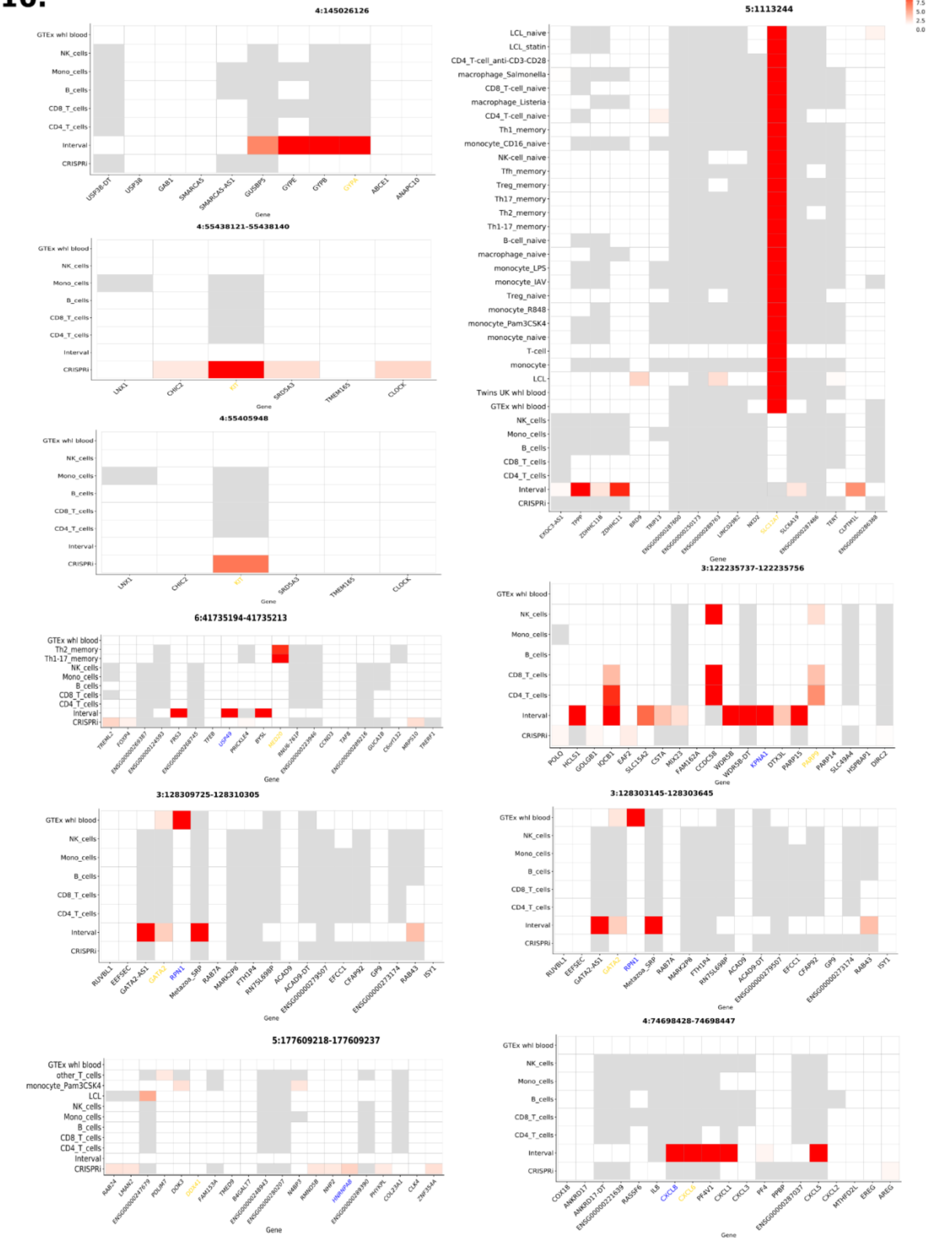

4:145026126
4:55438121-55438140
4:55405948
6:41735194-41735213
3:128309725-128310305
5:177609218-177609237
5:1113244
3:122235737-122235756
3:128303145-128303645
4:74698428-74698447
-log10 P-value
10.0
7.5
5.0
2.5
0.0
Gene

**11.**

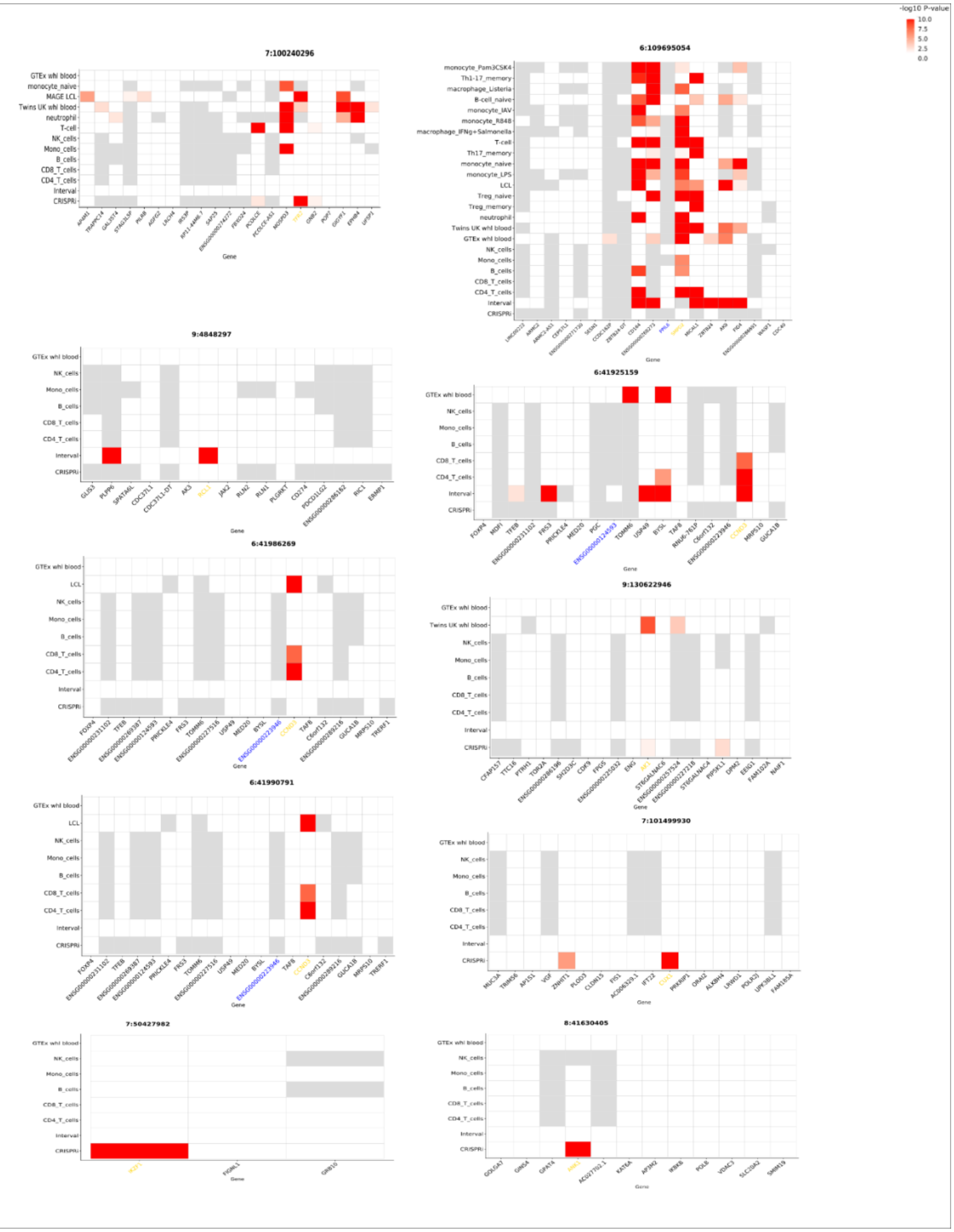
-log10 P-value
10.0
7.5
5.0
2.5
0.0
7:100240296
6:109695054
9:4848297
6:41925159
6:41986269
9:130622946
6:41990791
7:101499930
7:50427982
8:41630405
Gene

# 12.

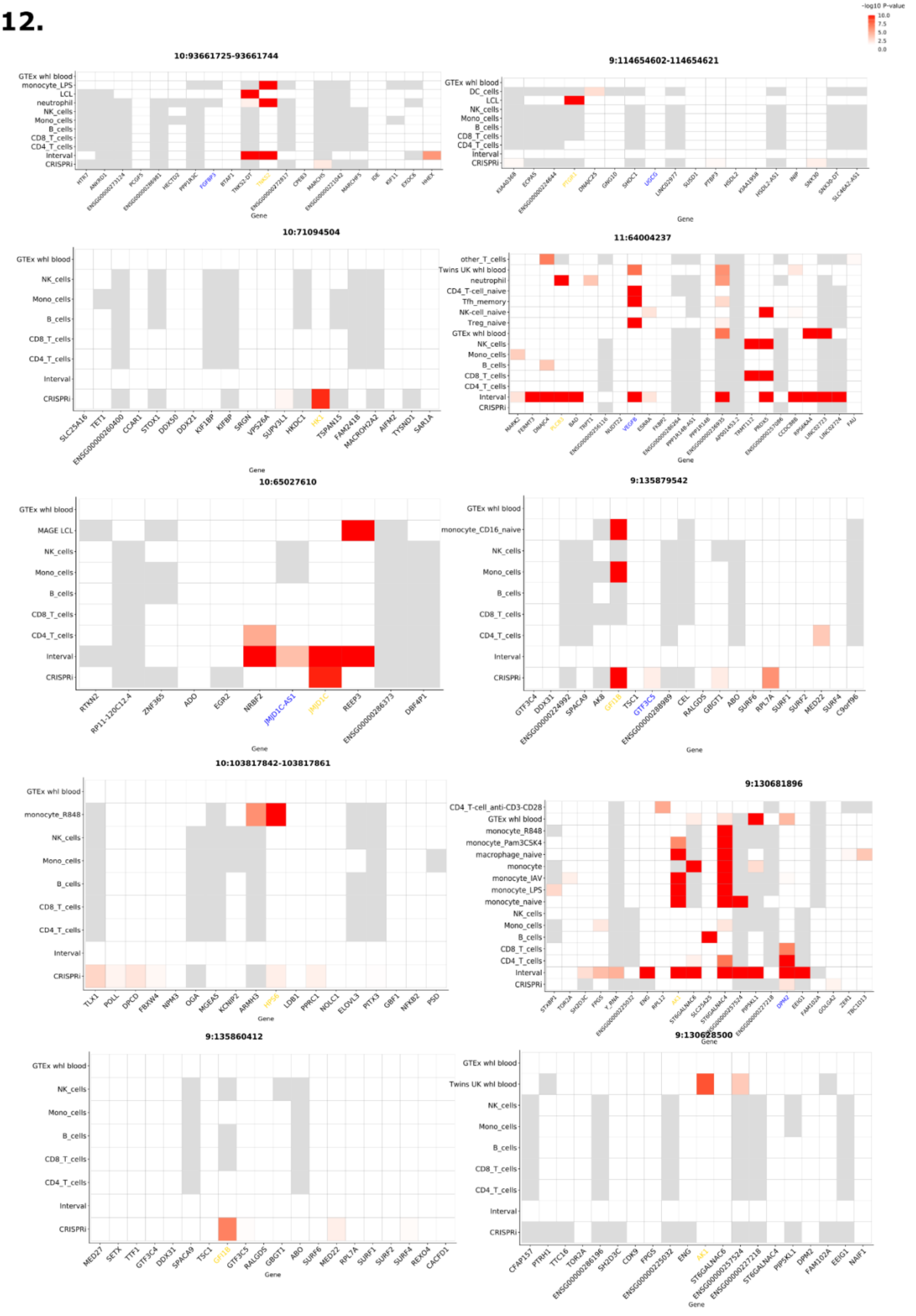
-log10 P-value
10.0
7.5
5.0
2.5
0.0
10:93661725-93661744
9:114654602-114654621
10:71094504
11:64004237
10:65027610
9:135879542
10:103817842-103817861
9:130681896
9:135860412
9:130628500
GTEx whl blood
Twins UK whl blood
NK_cells
Mono_cells
B_cells
CD8_T_cells
CD4_T_cells
Interval
CRISPRi
Gene

**13.**

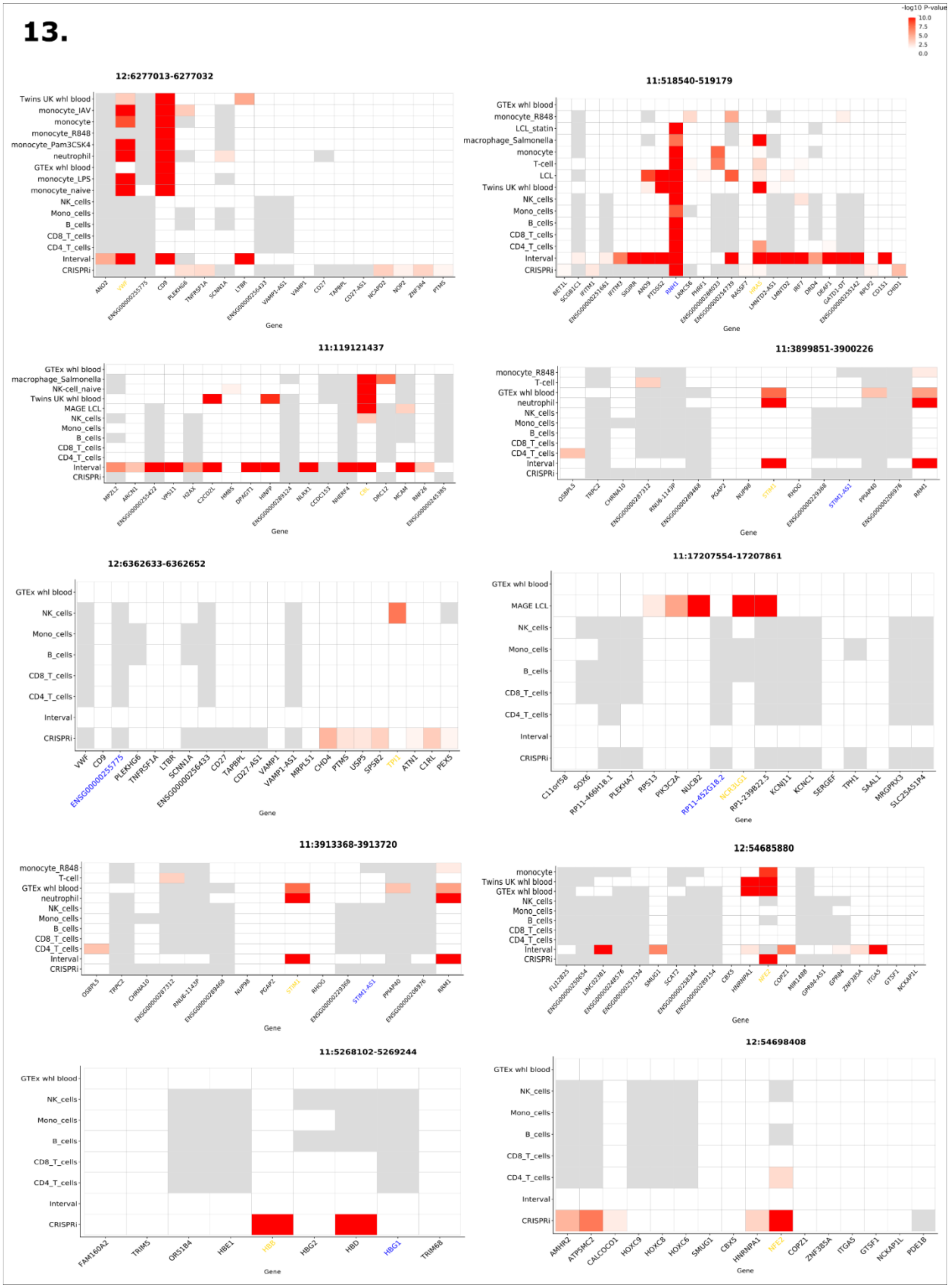
-log10 P-value
12:6277013-6277032
11:518540-519179
11:119121437
11:3899851-3900226
12:6362633-6362652
11:17207554-17207861
11:3913368-3913720
12:54685880
11:5268102-5269244
12:54698408
Gene

14.

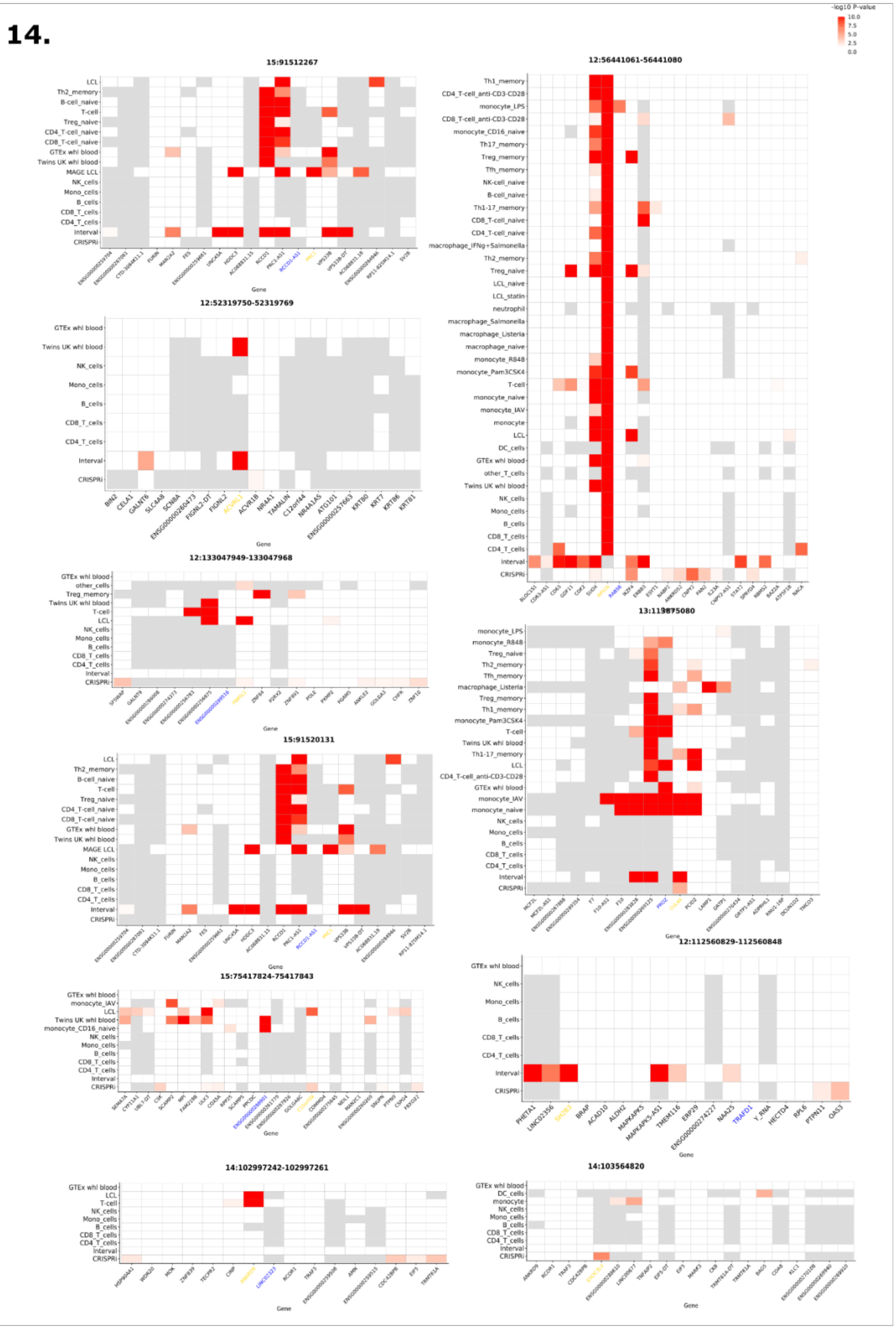

-log10 P-value
10.0
7.5
5.0
2.5
0.0
15:91512267
12:56441061-56441080
12:52319750-52319769
12:133047949-133047968
15:91520131
12:112560829-112560848
15:75417824-75417843
14:102997242-102997261
14:103564820
Gene

**15.**

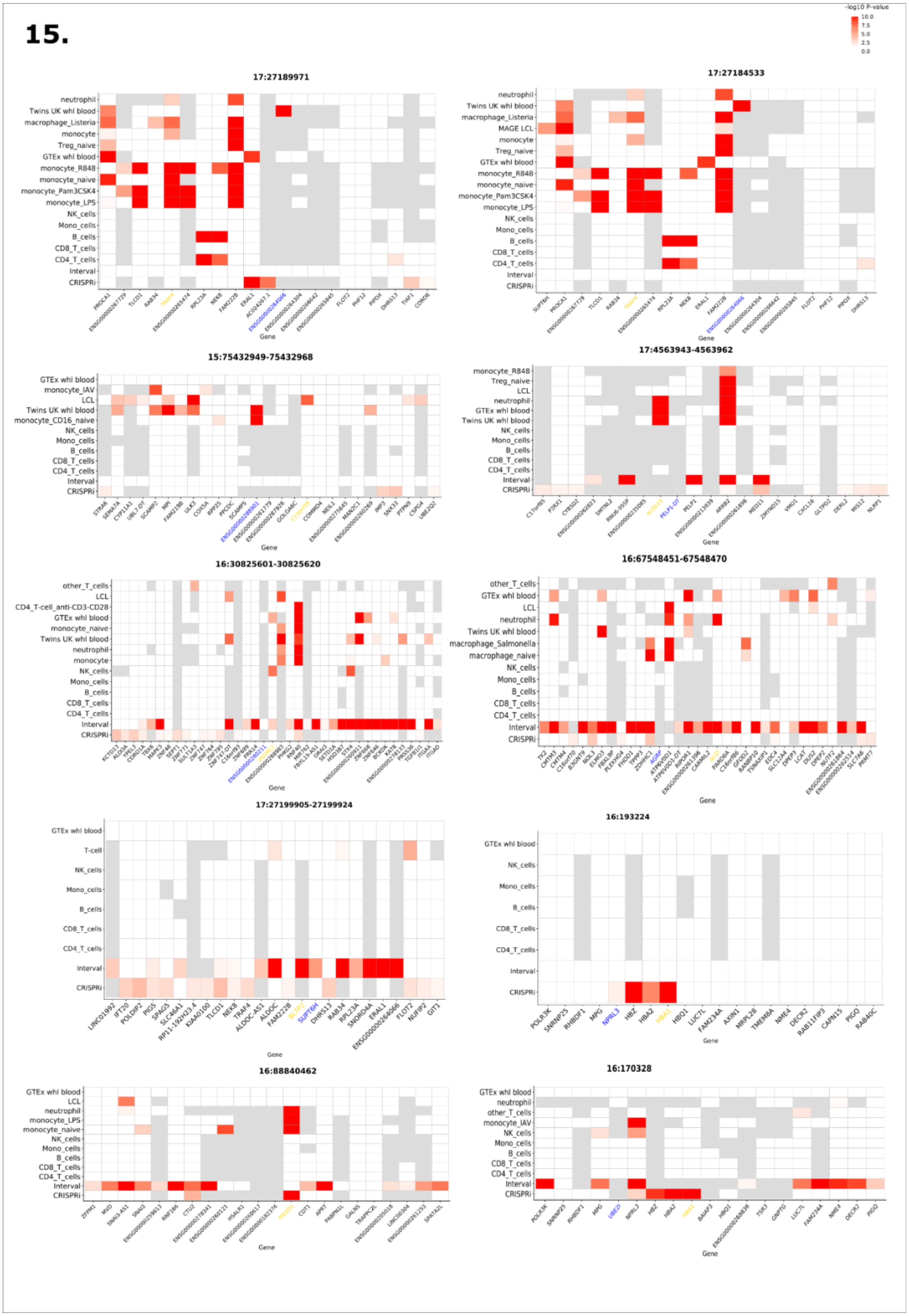
-log10 P-value
10.0
7.5
5.0
2.5
0.0
17:27189971
17:27184533
15:75432949-75432968
17:4563943-4563962
16:30825601-30825620
16:67548451-67548470
17:27199905-27199924
16:193224
16:88840462
16:170328
Gene

**16.**

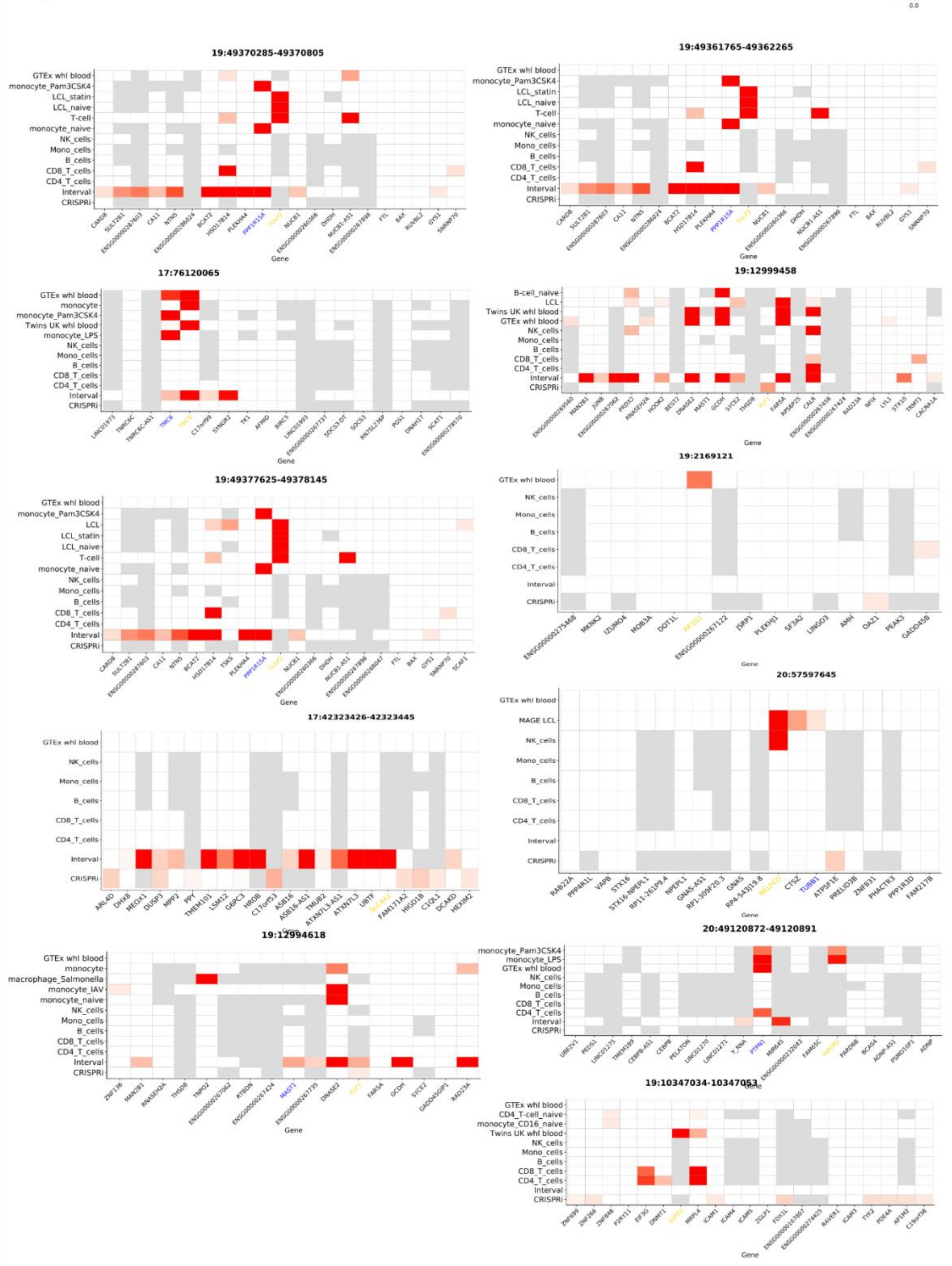
-log10 P-value
10.0
7.5
5.0
2.5
0.0
19:49370285-49370805
19:49361765-49362265
17:76120065
19:12999458
19:49377625-49378145
19:2169121
20:57597645
17:42323426-42323445
19:12994618
20:49120872-49120891
19:10347034-10347053
Gene
