## Supplementary Text for "CRISPRi perturbation screens and eQTLs provide complementary and distinct insights into GWAS target genes"

#### Defining regions for fine-mapping

To define fine-mapping regions, we first identified significant associations for each of the 29 blood cell traits from the UK biobank, selecting SNPs with p-values $< 6.6 \times 10^{-9}$. For each association, we defined the fine-mapping region as all SNPs within ±250 kb of the significant SNP. Overlapping regions were iteratively merged until no regions shared common SNPs, yielding a set of fine-mapping regions for each of the 29 blood cell traits. To create a unified set
across all traits, we further merged regions from different traits whenever they overlapped. This
process resulted in 899 disjoint fine-mapping regions, ensuring that each significant SNP ($p < 6.6 \times 10^{-9}$), and all SNPs within 250 kb of it belonged to exactly one region. The resulting fine-mapping regions varied in size, ranging from 500 kb to 13.47 Mb (mean: 1,325 kb; median:
949 kb). Notably, we excluded the extended MHC region (chr6: 25–36 Mb) due to its complexity
and challenges in interpretation, as it is typically analyzed separately.

#### Recalling eQTL data from OneK1K

#### Single-cell data collection

The OneK1K consortium is a research initiative that consists of single-cell RNA sequencing (scRNA-seq) data from 1.27 million peripheral blood mononuclear cells (PMBCs) collected from 982 donors of Northern European ancestry [1]. We downloaded the cell-by-gene data from the Human Cell Atlas (HCA).

https://cellxgene.cziscience.com/collections/dde06e0f-ab3b-46be-96a2-a8082383c4a1

We emailed the authors to request the raw VCF file, which contained data from 1,104 individuals and 759,993 markers.

#### Processing of genotypes

We used PLINK 2.0 to process the genotypes following the pipeline used in the original publication [2]. Briefly, we screen for bi-allelic markers, SNP missingness, and low genotyping of individuals

(SNP and individual call rates greater than 0.97), minimum MAF of 0.01, and Hardy-Weinberg equilibrium failure (p<10-6). We excluded samples with excess autosomal heterozygosity (± 3 SD from the mean). Then, we generated a genetic relationship matrix from all the autosomal SNPs using GCTA [3], and we removed any pair of individuals with an estimated relatedness larger than 0.125. After these quality control and processing steps, we kept 526,169 autosomal SNPs on chromosomes 1 to 22 from 1034 individuals. Finally, we performed imputation on chromosomes 1-22 using the Michigan Imputation Server with the Haplotype Reference Consortium panel (HRC r1.1 2016) [4] and ran using Minimac3 and Eagle v2.3 on Michigan Imputation Server (Loh et al. 2016). We kept SNPs with INFO > 0.8, R2 > 0.8, and MAF > 0.05. We generated a final catalog of 5,273,508 markers from 1,034 individuals.

### Cell classification

We obtained cell-type classifications by scPred [5] from the authors of the original paper (14 cell types; see more details in ref [1]). Additionally, we performed automated cell-type classification using Azimuth [6] to annotate the cells. Specifically, we mapped OneK1K query dataset to a previously published CITE-seq reference dataset of 162,000 PBMCs measured with 228 antibodies [7]. Following the publicly available tutorial, we used a set of specific functions from the Seurat R package v4.0.0 [6,8]. In brief, we normalized the reference dataset using the SCTransform function, we found anchors between reference and query, and we then transferred cell type labels and protein data from the reference to the query using the FindTransferAnchors function. We classified the OneK1K data using both the low-resolution cell-type-annotations (celltype.l1) and the high-resolution cell-type-annotations (celltype.l2).

### Identification of covariates influencing gene expression

*Population structure.* To remove any effect of population structure on gene expression, we obtained the genotype principal components (PCs) from the authors of the OneK1K paper [1]. We included the first six PCs as covariates in all our subsequent analyses.

*Hidden factors.* For each subpopulation and each cell-type classification method, we calculated the pseudo-bulk expression as the mean expression per gene per individual and then quantile-normalized and z-transformed. We then inferred latent variables using the recently published guidelines for pseudobulk eQTL studies (option 11) [9]. Briefly, we used the R package “peer” (v1.0) to generate the PEER factors (PFs) for the single-cell data, applying max iterations = 2000 and the number of PFs = 50. We used the R function *prcomp* with default parameters to calculate the first 50 Principal Components (PCs). We filtered out genes with a high proportion of zero expression across individuals ($\pi 0 >= 0.9$), we then log-transformed the gene expression data per gene ($\log(x + 1)$), and we finally standardized the data by applying z-score scaling per gene. To compare both estimates of hidden factors, we used the R package corrplot (v0.92) to calculate the correlation between the first 10 PFs and PCs. Since both estimates were highly similar, we decided to conduct our subsequent analysis using PCs. To choose an optimal number of latent variables fitted in the eQTL mapping, we used the elbow method from the R package PCAtools ("GitHub - kevinblighe/PCAtools: PCAtools: Everything Principal Components Analysis," n.d.).

**Single-cell cis-eQTL mapping**

The cis-eQTL association analysis was performed by tensorQTL v1.0.9 [10]. We used age, sex, the first 6 genotype PCs, and the expression PCs detected previously by the elbow method. We tested the SNPs located in a cis-window of ±1,000,000 around the TSS and with minor allele frequency > 5%. To be able to compare our cis-eQTL mapping method with the published catalog from the OneK1K consortium, we performed cis-eQTL mapping to predicted cell types from scPred, in addition to Levels 1 and 2 from Azimuth. We performed conditional analysis to discover independent cis-eQTLs with the tensorQTL mode *-cis_independent*. We considered significant cis-eQTLs with FDR < 0.05. Finally, to properly compare the power to detect cis-eQTLs in both studies (our study vs. the OneK1K study), we calculated the pi1 statistic for both OneK1K cis-eQTLs and our tensorQTL cis-eQTLs.

*TensorQTL pi1 = 1 - pi0 from TensorQTL p-values of OneK1K cis-eQTLs*

*OneK1K pi1 = 1 - pi0 from OneK1K p-values of TensorQTL cis-eQTLs*

**Multivariate Logistic Regression to Identify Features Associated with Shared Gene Targets**

To identify features associated with genes jointly identified by CRISPRi and eQTL methods, we performed a multivariate logistic regression analysis. Genes within 1 Mb of GWAS CREs were classified as either intersecting (identified by both methods) or non-intersecting (identified by only one). We excluded duplicate target genes across different GWAS/eQTL combinations and retained only significant cGenes and eGenes. For cGenes, gene expression was used to assess whether eGenes were expressed in K562 cells; for eGenes, expression across eQTL tissues and cell types was assessed using publicly available eQTL summary statistics from OneK1K, MAGE, GTEx, and the eQTL Catalogue.

We obtained gene-level metrics from multiple sources, including pLI, Episcore, the number of linked enhancers, and expression in GTEx whole blood. GWAS summary statistics and eQTL p-values were transformed to log-scale and averaged across loci per gene. Each gene was assigned a binary outcome variable indicating whether it was identified by both CRISPRi and eQTL methods (intersecting) or by only one (non-intersecting). Genes were also annotated as gold-standard if they overlapped with rare coding variant associations.

Logistic regression was performed using a binomial generalized linear model with the following predictors: scaled colocalization probability (PP.H4), Episcore, TSS distance to the CRE (scaled per 100 kb), gene expression (log-transformed median TPM), eQTL and GWAS significance (-log p-values), number of enhancers, and pLI status (≥0.9). Odds ratios and p-values were derived from the fitted model to assess the association between each feature and the likelihood of a gene being identified by both methods.

### Power calculations

#### Bulk eQTL Power Calculation

eQTL power is defined as the probability **P(i ∈ S)** that gene *i* belongs to the set of significantly differentially expressed genes *S*. This quantifies the statistical power (i.e., the probability of rejecting the null hypothesis $H_0$ when the alternative hypothesis $H_1$ is true) for detecting an eQTL at gene *i*. The calculation assumes an effect size of 0.5.

For genes with high expression levels (mean count > 5), eQTL power is estimated analytically using an F-test [11]. This power depends on sample size, the significance threshold α ($1 \times 10^{-3}$), and the proportion of variance explained ($q^2$) by the variant in the population. Assuming genotypes are in Hardy-Weinberg equilibrium, $q^2$ is given by:

$$q^2 = 2 \times MAF \times (1 - MAF) \times \beta^2$$ [12]

In this context, the F-test power estimate represents the probability of correctly rejecting $H_0$ when $H_1$ is true. For example, a power of 0.8 (80%) indicates an 80% chance of detecting a true eQTL effect. For genes with low expression counts (<5), analytical power estimates tend to be overly optimistic. Instead, power is estimated through simulations using scPower [11].

**Bulk eQTL Simulation-Based Power Calculation**

To determine simulation-based power for a given sample size $n$, significance threshold $\alpha$, and mean count $\mu c$ of the lower-expressed allele, the following steps are repeated $B = 100$ times:

1. **Genotype Simulation**: To account for the discrete nature of genotypes, allele frequency $f_a$ is drawn from a uniform distribution between 0.1 and 0.9.
2. **Read Count Simulation**: Given $f_a$, the effect size $\beta$, and the standard deviation of residuals, gene expression counts ($x$) are sampled from a negative binomial distribution parameterized for each genotype $g$.
3. **Statistical Testing**: A linear regression model is used to calculate the p-value for the null hypothesis ($H_0$: no eQTL effect).

The power is then determined as the proportion of simulations where $P < \alpha$ ($1 \times 10^{-3}$), reflecting the probability of correctly detecting an eQTL effect under these conditions.

**sc-eQTL Power Calculation**

**Expression Probability Calculation in scRNA-seq Experiments**

In scRNA-seq experiments, individual cells are typically not sequenced to saturation, resulting in sparse count matrices where only highly expressed genes are detected with nonzero counts. Additionally, both the overall number of transcripts and the expression levels of individual genes vary significantly across different cell types. To account for these factors in our power analysis, we estimated the expression probability of each gene using scPower [11]. Expression probability quantifies the probability $P(i \in E)$ that gene $i$ belongs to the set of expressed genes $E$. The overall power to detect a single-cell eQTL (*sc-eQTL*) for a given gene is then calculated as the product of eQTL power and expression probability.

**Modelling Expression Probability**

The total gene counts (*y*) across all cells follow a negative binomial distribution, with parameters adjusted based on the number of cells per cell type and per donor. The probability that the summed counts *y* exceed a given threshold *n* is expressed as:

$$p_{i,s} = P(y_{i,c,s} > n) = 1 - F_{NB}(n, \mu'_{i,c,s}, \phi'_{i,c,s})$$

A gene *i* is considered expressed in a given cell type c if the sum of counts *y* across all cells of that type within an individual *s* exceeds *6* in more than *20%* of individuals. The expression probability of gene *i* is then obtained using the cumulative binomial distribution (*FBin*), which accounts for the proportion of individuals with gene expression exceeding the threshold.

$$P(i \in E) = 1 - F_{Bin}(k \cdot n_s, n_s, p_{i,s})$$

Several factors influence the calculation of expression probability in scRNA-seq analyses. Sample size (n) determines the probability that a given percentage of individuals exhibit gene counts greater than *n*, affecting overall detectability. The number of cells (n cells) is used to model the summed negative binomial distribution, capturing how cell count variations impact expression probability. Cell type frequencies influence the probability that a proportion of individuals have gene expression levels exceeding the threshold $y > n$, accounting for differences in the representation of cell types within the dataset. Additionally, the percentage of individuals surpassing the count threshold *n* is directly incorporated into probability calculations.

**Intersection with ATAC-seq peaks**

We next assessed whether GWAS CREs overlapped regions of open chromatin in various hematopoietic lineages. Specifically, we used bedtools to intersect GWAS-implicated CREs with ATAC-seq peaks derived from B cell, $CD4^+$ T cell, $CD8^+$ T cell, Common Lymphoid Progenitor, Common Myeloid Progenitor, Erythroblast, Granulocyte-Monocyte Progenitor 1 (low), Granulocyte-Monocyte Progenitor 2 (mid), Granulocyte-Monocyte Progenitor 3 (high), Hematopoietic Stem Cell, Lymphoid-Primed Multipotent Progenitor, Megakaryocyte, Megakaryocyte-Erythroid Progenitor, Multipotent Progenitor, Monocyte, Natural Killer cell,

Myeloid Dendritic Cell, and Plasmacytoid Dendritic Cell [13]. Overlaps were defined by the direct intersection of variant coordinates with ATAC-seq peaks.

**Enrichment for gold-standard genes**

The overlap enrichment between cGenes/eGenes and gold-standard genes was assessed using random permutations of gene sets. The analysis followed these steps:

1. Randomly select a set of genes from the Ensembl v99 list of protein-coding genes using the R function sample, ensuring that the selected set matches the size of the test set.
2. Overlap the randomly selected gene set with gold-standard genes to determine how many of the randomly chosen genes overlap with the gold-standard set.
3. Repeat this process 10,000 times to generate a background distribution of expected overlaps under the null hypothesis.
4. Compare the observed overlap with this background distribution to assess statistical significance with a Fisher's exact test.
